# Reference-Based Library Construction Improves Performance in Low-input Workflows

**DOI:** 10.64898/2026.04.29.721088

**Authors:** Joshua Charkow, Mahmoudreza Ghaznavi, Brendon Seale, Jiaxi Peng, Anne-Claude Gingras, Hannes L. Röst

## Abstract

In low input mass spectrometry-based proteomics, Data Independent Acquisition (DIA), is quickly becoming the method of choice for label free quantification. Whether using empirical or *in silico* spectral libraries, performance is dependent on the library; however, the optimal library construction strategy for low input proteomics remains an open question. To address this, we examine and develop library construction approaches that are compatible with both spectrum-centric and peptide-centric analysis workflows. These approaches leverage a closely related, high-quality sample to improve library quality. First, we validated our approach in bulk sample amounts where we observed that the effects of gas-phase fractionation based library construction is dependent on the software framework, with improvements more pronounced in OpenSWATH compared to DIA-NN. In OpenSWATH, our peptide-centric library reconstruction workflow consistently outperforms a transfer learning strategy, an emerging alternative approach. In DIA-NN, trends are dependent on library source highlighting OpenSWATH’s stronger dependence on the search space. In low-input applications, such as single-cell-equivalent injection amounts (100 pg) of HeLa cell digest on a timsTOF SCP, our library construction approach provided more pronounced improvements across both software tools compared to bulk samples. Using a peptide-centric reconstruction approach with the OpenSWATH analysis framework, we detected over 15,000 peptide precursors (2480 protein groups), a 90% improvement over the original library. Furthermore, using a spectrum-centric construction approach, peptide precursor identification rates improved over 6-fold ( ∼1000 to ∼6000). Our strategy provides a practical solution for generating high-quality libraries in low-input applications.

## Introduction

Given recent technological advances, mass spectrometry-based proteomics is becoming increasingly capable of analyzing nanograms to picograms of peptide mass, enabling applications in single cell proteomics ^1,2^ laser capture microdissections,^3^ and enrichments from limited samples.^4^ For label free quantification in low input applications, Data Independent Acquisition (DIA) is growing in popularity because it achieves high proteome coverage in applications where sample amounts are limited.^5–7^ However, DIA multiplexes MS2 spectra which may result in high spectral complexity. This means that the traditional spectrum-centric analysis tools, which assume one precursor per spectrum, may not be directly applicable.

To mitigate the high degree of spectral complexity, a peptide-centric analysis workflow can be used for identification and quantification of peptides in DIA. This approach requires a peptide library containing *a priori* knowledge on peptide precursors to search for in the sample, and their peptide properties including mass-to-charge (m/z) and retention time to determine where the precursor’s signals are expected to be found. For each precursor, peptide properties are used to computationally extract signals from the multiplexed spectra resulting in a series of chromatograms. These chromatograms can be used to determine if the signals adequately match the library and for quantification.^8^ Given this strategy’s strong dependence on the library, the importance of library quality is widely recognized in the community. Many studies focus on the effects of using different libraries in analysis ^9–12^ and also attempt to systematically quantify the library’s quality.^13^ In general, a high quality library is regarded as one that maximizes proteome coverage, reproducibility and quantification accuracy.^9–12^

Offline fractionation of experiment-specific samples has long been considered the gold standard for generating high-quality peptide libraries,^14^ but this is impractical in low input sample amounts, such as in single cell proteomics. Currently, it is still an open question how to best construct a library for single cell proteomics applications. Some library generation strategies that are currently being explored include generating an experimental library from bulk samples,^6,7^ a reference library,^15^ an *in silico* library,^16,17^ or constructing the library directly from the data.^18^ However no single strategy has been universally adopted. Consequently, ad hoc library construction approaches are often used in single cell proteomics which may vastly affect proteome coverage, reproducibility and quantification accuracy. To minimize the effects of poor library quality, match-between-run (MBR) strategies are often employed,^19–21^ with some approaches extending the matching to high input reference samples.^22–25^ However, this inflates false discovery rates (FDR) in heterogeneous bulk samples,^26^ and heterogeneous single cells.^27^

Alternatively, the effects of poor library construction can be minimized by using a closely related, high quality reference sample to generate high quality libraries. This approach was pioneered by Searle et al., 2018 who found that using gas-phase fractionated (GPF) samples could refine experimental,^28^ reference ^28^ and *in silico* ^29^ libraries, resulting in increased proteome coverage and reproducibility. Although variations of this approach are starting to be adapted for single cell and small sample amount proteomics,^20,21^ key questions remain. This includes which reference samples work best for small sample amounts, how to incorporate information from reference samples, and the applicability of this approach across different experimental, *in silico* and reference libraries.

Here, we propose a library construction strategy for small sample amounts which uses a reference sample to improve library quality. This approach can be adapted for both peptide-centric and spectrum-centric analysis strategies. First, we validate this in bulk sample amounts across various initial libraries and software tools. Next, we adapt this strategy for small sample amounts where we use a reference sample of a higher protein mass in lieu of the GPF samples. Applying our approach in a small sample regime of 100 pg allowed us to improve peptide precursor identifications by 6-fold using a spectrum-centric approach with diaTracer and improve precursor identifications in a peptide-centric context almost 2-fold using OpenSWATH.

### Experimental Procedure

#### Experimental Design and Statistical Rationale

This study only consisted of technical replicates which are defined as repeat injections of the same digest. No biological replicates were used since all samples were sourced from commercial digests. For K562 bulk digests, 3 replicates were used consistent with standard practice. For the K562 dilution series, only a single replicate was used per dilution as this analysis was exploratory in nature. When applying our methods to single-cell equivalent injections we increased the number of technical replicates to 9 to account for the larger variation expected from smaller sample amounts.

For all performance metrics, the reference-based libraries were compared against an initial library as a baseline. Comparisons between the initial and reference-based libraries are descriptive and no statistical hypothesis tests were applied.

#### K562 Data Acquisition

K562 lysate was purchased from Promega (V651) and diluted to appropriate loading amounts (250 ng to 1 ng) in 0.1% formic acid. K562 samples were acquired using a self-packed and pulled silica column coupled to a timsTOF Pro 2. The column was packed with 1.9 µm ReproSil-Gold 120 C18 resin (Dr. Maisch) and maintained at 50 °C. Mobile phases A and B were 0.1% formic acid in water and 0.1% formic acid in acetonitrile, respectively. Samples were acquired on a 90 minute active gradient ramping the fraction of B from 5% B to 35% followed by 15 minutes at 85% B. Peptides were separated at a flow rate of 100 nL/min. Chromatography was coupled to mass spectrometry using a 1600 V electrospray. All runs used an accumulation and ramp time of 100 ms. For the standard DIA runs, the scheme consisted of 25 m/z windows ranging from 400–1200 m/z and 0.73 to 1.46 in mobility (1/k_0_) with 4 mobility ramps as published in Meier et al., 2020.^30^ Combined with the MS1 scan this resulted in a total cycle time of 1.7s. Collision energy was selected based on mobility ranging from 21 ev at 0.8 1/k_0_ to 65 eV at 1.5 1/k_0_. The GPF scheme consisted of 13 injections with windows ranging from 400–1200 m/z and a mobility range of 0.6 to 1.60 1/k_0_ (Table S1). Each injection covered a 60 m/z region using 5 m/z isolation windows with a 1 m/z overlap between windows. The last two injections covered an 80 m/z region. Combined with the MS1 scan this resulted in a total cycle time of 1.4s. Injections contained one overlapping window with the previous injection.

#### Single-Cell Equivalent Data Acquisition

Pierce HeLa lysate (# 88329) was diluted to appropriate loading amounts with 0.1% formic acid and 0.2% n-dodecyl-beta-D-maltoside. Samples were acquired on an EvoSep One LC coupled to a timsTOF SCP. Samples were loaded onto the EvoSep EvoTip Pure using the standard procedure for analysis. Peptides were separated using the EvoSep One LC on a standard EvoSep Whisper 40 SPD gradient coupled with the IonOptics Aurora 3 Elite (15 cm x 75 μm). DIA runs used a custom scheme with m/z windows ranging from 400–1000 m/z, a mobility range of 0.64–1.45 1/k0 and 3 mobility ramps (Table S2). Combined with the MS1 scan this resulted in a total cycle time of 0.9s. Collision energy was ramped according to mobility increasing from 20 eV at 0.6 1/k_0_ to 59 eV at 1.6 1/k_0_.

#### K562 Experimental Library Generation

Promega K562 (V6951) lysate was fractionated according to the manufacturer’s instructions for Pierce High pH Reversed-Phase peptide Fractionation Kit (# 84868) with a 100 µg input and fractionations at 5%, 7.5%, 10%, 12.5%, 15%, 17.5%, 20%, 50% acetonitrile. Samples were acquired on a timsTOF Pro 2 as outlined above. Each fraction was injected into a timsTOF Pro 2 acquired using ddaPASEF. To generate the experimental library, we analyzed the high pH reverse phase fractionation injections acquired in Data Dependent Acquisition (DDA) using Fragpipe (v22.0) which includes Philosopher (v5.1.1),^31^ MSFragger (v4.1),^32–34^ MSBooster (v1.2.31),^35^ Percolator (v3.06.5),^36^ Protein Prophet (v5.1.1)^37^ and Easypqp (v.0.1.50).^34^ Minor changes from the default parameters (which includes a protease of stricttrypsin, a fixed modification of carbamidomethylation on cysteine, a variable modification of oxidation on methionine, a variable modification of acetylation on the N-terminus, a fragment tolerance +/-20ppm, and up to 2 missed cleavages) were made for library generation and mainly to decrease the search space to be more in line with the *in silico* library. Changes include deactivating de-isotoping, setting the maximum precursor tolerance +/-50 ppm, variable modifications per peptide to 1, the maximum digest length to 30, and minimum matched fragments to 6. The spectra were searched against the human proteome (UP000005640) downloaded on 12/09/2024 and common contaminants added by Fragpipe. This library was filtered to a 1% PSM, protein and peptide level FDR.

#### *in silico* Library Generation

The *in silico* library was generated using alphapeptdeep (v1.3.0)^38^ with the human reference proteome and contaminants described above. Digestion was done using the “trypsin/P” enzyme with the max missed cleaves to 1, carbamidomethyl fixed modification, peptide length from 7–30 and precursor charges from 2+ – 4+. The built in timsTOF models were used for peptide property prediction. Fragment ions were generated with charge states up to 2+.

#### Published Libraries

The PanHuman library was downloaded from Rosenberger et al., 2014 (PXD000953).^39^ Ion mobility predictions were appended using the alphapeptdeep models.^38^ For consistency with the other libraries, the protein annotations were re-annotated with the previously downloaded reference proteome using DIA-NN, and decoys were removed. The timsTOF library was downloaded from Ctortecka et al., 2024 (MSV000093867).^15^

#### Direct Library

The direct libraries were created by analyzing the DIA runs directly using diaTracer (v1.1.5)^18^ in Fragpipe. Search settings are as described above except that the maximum precursor tolerance was set to +/-10 ppm as recommended by FragPipe.

#### Analysis and Library Reconstruction With OpenSWATH-PyProphet Workflow

Analysis was conducted using OpenMS (v3.4) ^30,40^ and Pyprophet (v3.0.2).^41,42^ First, libraries were processed with OpenSWATHAssayGenerator using the default parameters. For reconstructed libraries, settings were relaxed such that peptide precursors could contain 4–6 fragment ions since these libraries are expected to contain a large proportion of true positives. Decoys were appended with OpenSWATHDecoyGenerator using the “pseudo-reverse” method and setting “-switchKR” to true. For OpenSWATHWorkflow, settings are as described in Meier et al., 2020^30^ with the additional “-pasef” flag enabled. For the *in silico* library “Scoring:stop_report_after_feature” was set to 2 to limit the output size. iRT files were generated using high confidence peptides detected by diaTracer. Pyprophet was run with the default parameters using the XGBoost classifier except when overfitting occurred in which case the support vector machine classifier or linear discriminant analysis was used. Overfitting was determined by manual examination of the target-decoy distribution using the output PDF files provided by Pyprophet to ensure that the false target distribution was representative of the decoy distribution. For the direct library in small sample amounts, the λ value in π_0_ estimation was decreased to 0.001 on the peptide and protein level to ensure π_0_ estimation would not fail. In the cases where reference sample(s) were analyzed, the reconstructed library was exported using the new “pyprophet library” command with the default parameters. Peptide precursors were quantified by summing up the area intensities of all fragment ions (MS2 level), as is standard in OpenSWATH.

#### Analysis and Library Reconstruction With DIA-NN

Analysis was conducted using DIA-NN (v1.9.2)^43,44^ on Linux using the following additional flags: “direct-quant” and “report-lib-info”. The parameters “mass-acc” and “mass-acc-ms1” were set to 15 ppm as recommended by the software. MBR was not used to allow the effects of library reconstruction to be evaluated independently from it. Library reconstruction used a custom script (accessible via https://github.com/jcharkow/reference-based-libraries). Briefly, this script aims to create the same library as DIA-NN would export internally with the “full-profiling” and “gen-spec-lib” flags; however, it also corrects for fragment ion intensity based on the experimental library and it filters entries at the protein 1% false discovery rate (FDR) level as well. Peptide precursors were quantified by summing up the area intensities of all fragment ions to be more comparable with OpenSWATH.

#### Transfer Learning

Transfer learning was conducted with alphapeptdeep.^38^ This involves supplying the reconstructed library to refine the default models and then regenerating the *in silico* library using the new models. For OpenSWATH, a modified version of the reconstructed library was supplied which contained more fragment ions per peptide so that fragment ion intensity transfer learning did not overfit due to a lack of fragment ions.

#### Chromatogram Visualization

Chromatogram visualization was conducted using massDash (v0.1.1)^45^ and pyopenms_viz (v.1.0.0).^46^ Scripts for analysis and figure generation are available at https://github.com/jcharkow/reference-based-libraries.)

## Results

### The Reference-based Library Construction Workflow

Our approach uses additional reference injections acquired in DIA that are closely related to the target sample in library construction and reconstruction. These injections are of higher quality (i.e. more protein mass injected, fractionated samples) than the target injection.

A high-quality library closely matches the experimental sample, which we assess using three metrics: (1) peptide property precision — the root mean square deviation (RMSD) in the property between the sample and the library, defined as

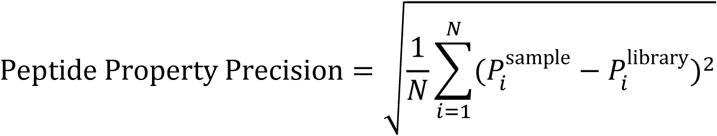

where *_P_* denotes a peptide property (i.e. RT, IM, relative fragment ion intensity) and *_N_* refers to the number of confidently identified peptide precursors (2) library sample concordance — the Jaccard index between the set of peptide precursors in the library and those in the sample, defined as

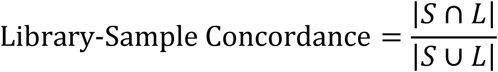

where *_S_* and *_L_* refer to the set of peptide precursors in the sample and library respectively and (3) sample scope — the total number of peptide precursors contained in both the sample and the library defined as

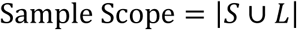

High-quality libraries typically require large amounts of the sample of interest (e.g. to be used in offline fractionation) to construct; however, this library construction strategy is unfeasible when sample amounts are limited.

Here, we present a workflow for generating high-quality libraries that can be applied with smaller amounts of additional material. Our library construction workflow comes in two different variations: peptide-centric library reconstruction and DIA-based spectrum-centric library construction. The peptide-centric reconstruction approach begins by analyzing the reference sample against an initial library. The results are then used to empirically correct peptide properties and remove undetected peptides, producing a higher-quality library (Figure 1). The spectrum-centric approach constructs a library directly from the DIA acquired reference sample rather than from the target sample itself (Figure S1). This results in a more comprehensive library compared to one directly generated from the target sample. In both approaches, the reference sample is used to create a high-quality library for peptide-centric analysis.

**Figure 1:**
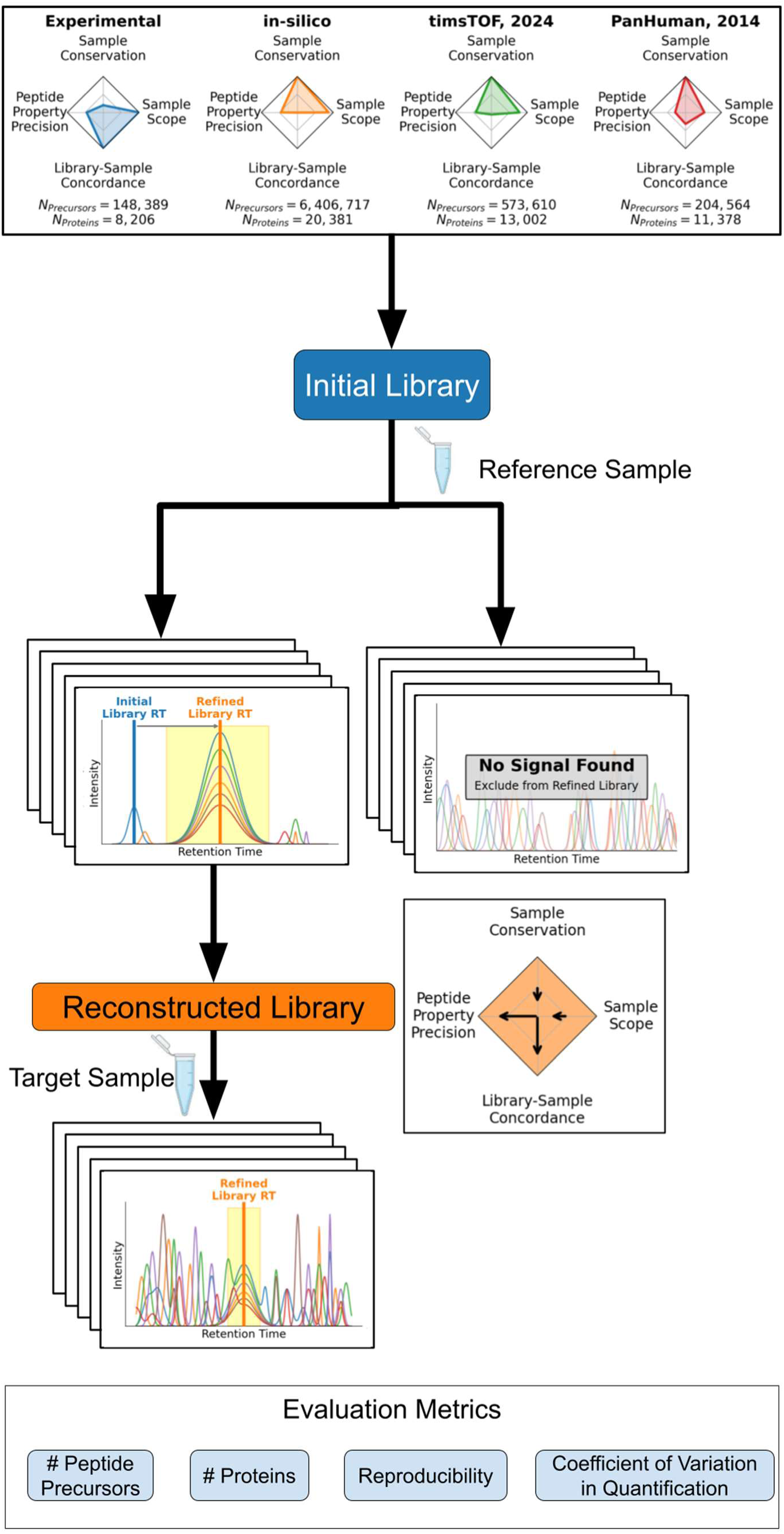
Overview of the peptide-centric library reconstruction workflow. Schematic of the peptide-centric library reconstruction workflow. Peptide libraries consist of a list of peptides and their corresponding characteristics (e.g. m/z, retention time, ion mobility). Different peptide libraries contain distinct trade-offs in terms of library quality and sample usage efficiency. Quality metrics depicted here include (1) sample scope — the proportion of peptides in the sample that are present in the library; (2) library-sample concordance — the similarity between the peptides in the library and the peptides in the sample; and (3) peptide property precision — the RMSD of retention time, ion mobility, and fragment ion intensities. Further information on the initial libraries tested can be found in Table 1. Only the experimental library requires additional sample specific injections and thus has the lowest sample conservation. The initial library is analyzed with the reference sample, and these results are used to reconstruct the library. Library reconstruction involves excluding peptide precursors not identified in the reference sample and correcting peptide characteristics based on the results of the reference sample. This increases peptide property precision and library-sample concordance at a slight cost in sample conservation and sample scope. The target sample is analyzed with the reconstructed library whose higher quality enables noisy peptide signals to be detected with higher confidence. The effects of library reconstruction are evaluated by comparing the number of peptide precursors, number of protein groups, reproducibility and the CV in intensity between the initial and reconstructed libraries. Created with BioRender.com (2026).

**Table 1:** Summary of Different Libraries.

| Library Name | Source | # Peptide Precursors | # Proteins |
| --- | --- | --- | --- |
| Experimental | Constructed from offline fractionated samples in DDA (see methods). | 148,389 | 8,206 |
| *in silico* | *in silico* digest the entire human proteome. Peptide characteristics are predicted by alphapeptdeep<sup>38</sup> | 6,406,717 | 20,381 |
| PanHuman, 2014 | Rosenberger et al., 2014.<sup>39</sup> | 204,564 | 11,378 |
| timsTOF, 2024 | Cell line reference library acquired on a timsTOF family instrument from Ctortecka et al., 2024.<sup>15</sup> | 573,610 | 13,002 |
| Direct | Constructed directly from target diaPASEF samples using diaTracer,<sup>18</sup> a spectrum-centric approach. | 70,132 | 6,822 |

A consequence of using a high-quality library is that the majority of false library entries are particularly difficult to distinguish from true signals, meaning that the reported FDR may underestimate the true rate of false identifications. To minimize the transferring of false entries, we ensured that all entries of the new library were identified at 1% FDR for precursor, peptide (if applicable) and protein-levels in the reference sample and precursors were also removed if they contained fewer than 4 transitions.

### Effects of Library Construction Using GPF is Dependent on Library Source and Software Tool

First, we validated our library construction workflow in bulk sample amounts by using reference samples of higher selectivity. The reference samples employed gas-phase fractionation (GPF) consisting of 13 injections covering the m/z range of 400–1200 (Figure S2, Table S1).

This acquisition scheme achieves higher selectivity compared with the standard scheme because isolation windows were only 5 m/z rather than 25 m/z wide. Here we employed a simple sampling scheme for GPF which only uses one mobility ramp time; however, more complex sampling schemes can be used to minimize the number of injections required for construction.^47^ GPF-based library construction results in a higher quality library increasing the confidence that the detected peak group can be confidently assigned to the peptide query (Figure 2A). We tested our strategy against various libraries that differed in quality, size, sample source and instrumentation used for acquisition (Figure 1, Table 1). Libraries tested include an offline-fractionated experimental library, two published reference libraries,^15,39^ an *in silico* library, and a library constructed directly from the target sample (spectrum-centric approach). When analyzing the GPF runs using the initial libraries, we did not observe a drop in precursor identification near the edges of the m/z isolation windows (Figure S3).

**Figure 2:**
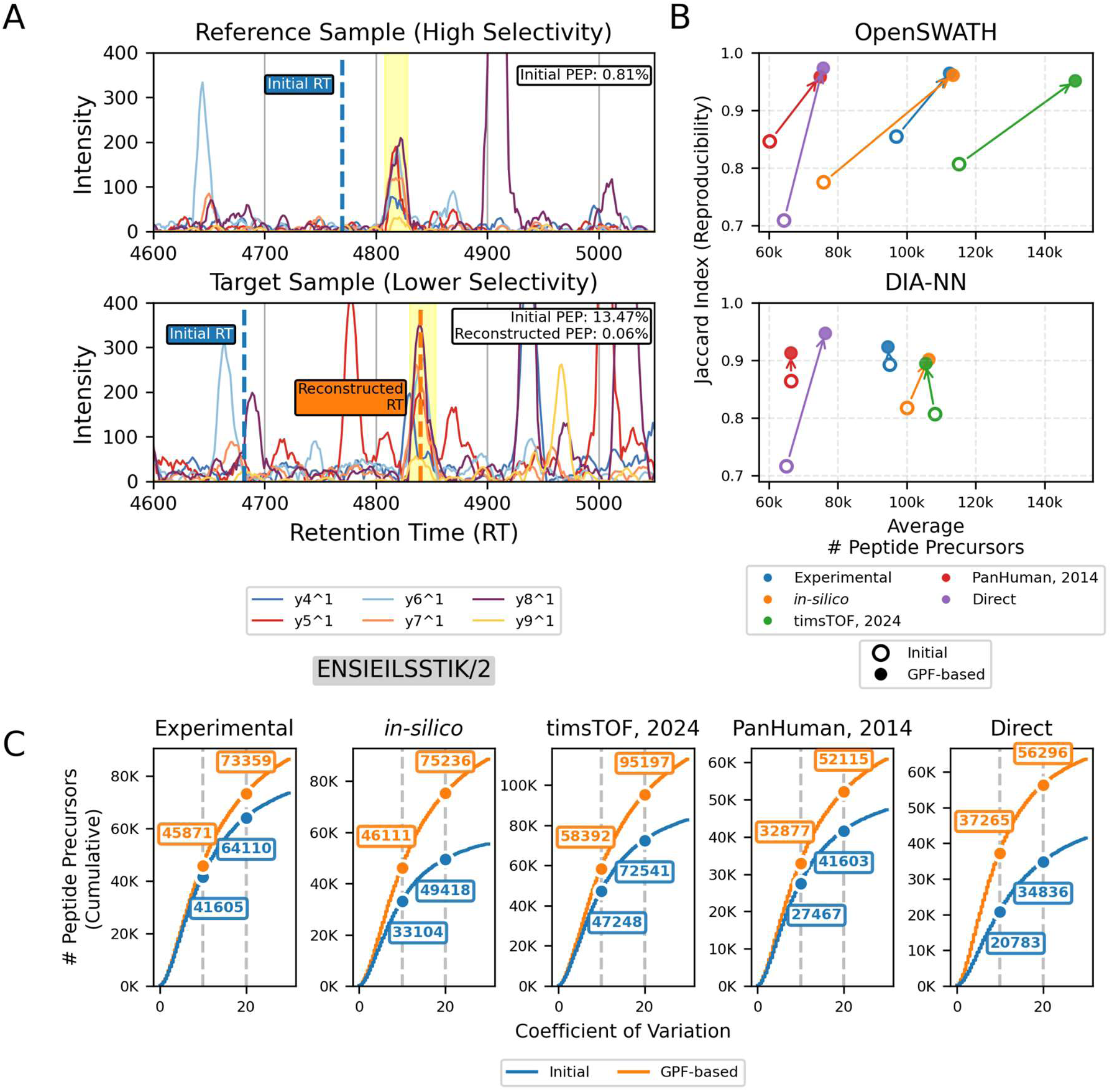
Impact of GPF-based library construction in bulk sample loads depends on software framework. (A) Example of a peptide precursor that is only identifiable after peptide-centric library reconstruction. A confident identification occurs when fragment ion traces belonging to the peptide of interest (represented here as lineplots) coelute with one another. Searching for peptide precursor *ENSIEILSSTIK^2+^*with the initial *in silico* library only yields a confident identification in the reference sample (Posterior Error Probability (PEP) = 0.81%) and not in the target sample (PEP = 13.47%). However, if the reconstructed library is used, this precursor can be confidently identified in the target sample (PEP = 0.06%). Note that the initial retention times occur at different locations even though the library is the same because of internal nonlinear retention time calibration. (B) Average number of peptide precursors identified across the three replicates against overall peptide precursor level reproducibility (Jaccard index) across libraries for two software tools: OpenSWATH (top) and DIA-NN (bottom). The results from the initial library are represented by a hollow circle, and the results from the GPF-based library are colored with a filled-in circle. Different colours represent the different libraries tested. Arrows compare the relationship between the initial and GPF-reference-based library for each library source. Arrows consistently point upwards indicating an improvement in reproducibility. For OpenSWATH, arrows specifically point to the upper right indicating improvements in precursor identifications and reproducibility. Trends are more pronounced in OpenSWATH compared to DIA-NN. Specifically, OpenSWATH exhibits 8-fold greater improvements in identifications and 1.7-fold improvements in peptide precursor level Jaccard index. Note that not all libraries share the same search spaces leading to differences in the maximum achievable number of peptide precursors. (C) Cumulative distribution of commonly identified peptide precursors (1% FDR) ranked by their CV in intensity on OpenSWATH results for the initial (blue) and GPF-based (orange) libraries. The GPF-based libraries quantify on average 10K and 18K more precursors with an intensity CV less than 10% and 20% respectively across all library sources.

To evaluate the effect of library quality, we compared the results from the three K562 digest technical replicates across the initial and the GPF-based versions of each library for both DIA-NN and OpenSWATH. Evaluation metrics include the number of peptide precursors identified, the number protein groups identified, precursor level reproducibility (summarized by the Jaccard index), and the number of confidently identified precursors with high quantitative precision (precursor-level intensity coefficient of variation (CV) < 20%). Across both software tools, GPF-based libraries always achieved higher reproducibility across the replicates (Figure 2B, Figure S4). Notably, OpenSWATH showed concurrent gains in identifications, improving both reproducibility and coverage simultaneously (Figure 2B, Figure S4, top panels). Although this trend was observed in DIA-NN for libraries with proteome wide-search spaces, incorporating GPF samples into library construction had a minimal impact on the number of identifications in libraries with smaller search spaces (Figure 2B, Figure S4, bottom panels). Overall, this demonstrates that the impact of reference-based library construction is more pronounced in OpenSWATH compared to DIA-NN. In OpenSWATH, even minimal improvements to peptide-property precision and library-sample concordance translate into an increase identifications and reproducibility.

Compared to the initial libraries, GPF-based libraries always increased the number of quantitatively precise peptide precursors across both software tools (Figure 2C, Figure S5). Since the shape of the cumulative distributions only visibly differ at their tails, this suggests that the gains in identifications of high-quality libraries primarily occur in noisier signals. However, it remained unclear whether these new precursor identifications were just due to gains in statistical power or whether the high-quality libraries affected the peak group chosen. To assess this, we took advantage of the fact that OpenSWATH performs greedy peak group detection outputting a peak group regardless of if a confident identification was made. We found that new common identifications resulting from library reconstruction, on average, had a lower CV compared to the initial library (Figure S6). This suggests that updating the peptide properties in peptide-centric library reconstruction corrects for errors in the peak group selected, achieving effects similar to MBR strategies.

GPF-based library construction consistently had the most pronounced effects on libraries with a proteome-wide search space, which include the direct library generated by the spectrum-centric approach and the *in silico* library. Our spectrum-centric library construction approach increased the average number of peptide precursors by over 17% (∼65K to ∼76K) and the Jaccard index by over 32% (∼0.7 to ∼0.95) for both software tools. The number of quantitatively precise precursors with a CV less than 20% improved by 87% (∼34K to 64K) in DIA-NN and 62% (∼34K to ∼56K) in OpenSWATH.

Similarly, peptide-centric reconstruction of the *in silico* library also increased identifications and reproducibility across both software tools. Using OpenSWATH, *in silico* library reconstruction increased the average number of peptide precursors by nearly 50% (∼75K to ∼113K), the Jaccard index by 24% (0.77 to 0.96), and the number of peptide precursors quantified with a CV of less than 20% by nearly 50% (∼49K to ∼75K). Using DIA-NN, trends showed improvement although less pronounced; peptide precursor identifications improved by 6% (∼100K to ∼106K), Jaccard index improved 10% (0.82 to 0.90), and the number of precursors quantified with a CV of less than 20% improved by 14% (∼74K to ∼84K) (Figure 2, Figure S5). Notably, this demonstrates that although OpenSWATH is poorly optimized for libraries with a large search space, library reconstruction can mitigate this by yielding superior reproducibility and precursor identification counts to DIA-NN before and after library reconstruction. The consistently pronounced improvements in the *in silico* compared with other peptide-centric library reconstructions can likely be attributed to the large improvement in library-sample concordance. In this context, reconstruction removes over 98% of precursors from the *in silico* library, drastically reducing the number of hypotheses (Figure S7).

The effect of incorporating GPF runs into library construction is more pronounced for OpenSWATH compared to DIA-NN. This indicates that OpenSWATH has a stronger dependency on library quality compared to DIA-NN. Nevertheless, when a very low-quality library is used such as an *in silico* library which has very low library sample concordance, strong improvements are observed across both software tools.

### Comparison Between Peptide-Centric Library Reconstruction and Transfer Learning Approaches

Another popular approach for refining libraries is transfer learning. In this approach, the *in silico* model is retrained on experimental data to provide more accurate predictions of peptide properties for the specific experiment being analyzed. Unlike the reconstruction approach presented above, transfer learning does not filter library precursors maintaining the sample scope of the library allowing for potentially greater proteome coverage. Yet this comes at the cost of the low statistical power associated with multiple hypothesis testing that is relieved by the filtering in our library reconstruction approach. Comparing transfer learning with our library reconstruction approach can provide insight into the trade-off between sample scope and library-sample concordance. Our library reconstruction approach prioritizes library-sample concordance while transfer learning prioritizes sample scope.

Here, transfer learning involves fine tuning the base models from alphapeptdeep^38^ using peptide precursors confidently detected in the GPF reference samples and using these new models to predict the peptide properties for all library peptide precursors. These new libraries resulting from transfer learning were then used to analyze the full DIA triplicates.

Our library reconstruction strategy consistently achieves higher reproducibility than the transfer learning strategy (Figure 3A), suggesting that a higher library-sample concordance is associated with increased reproducibility. This is further supported by the observations that library specificity accounts for 87% of reproducibility variance in OpenSWATH and 64% in DIA-NN (Figure S8).

**Figure 3:**
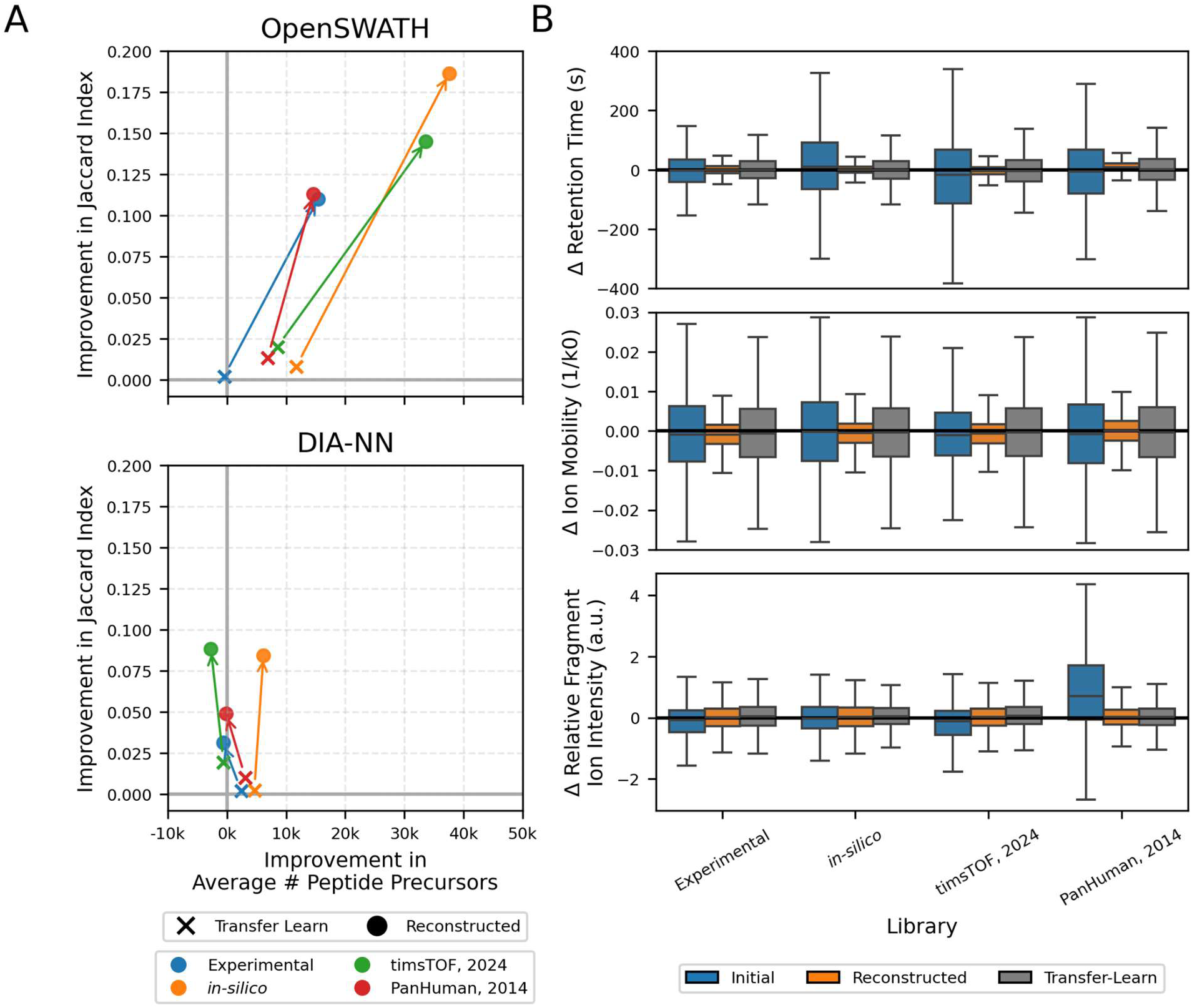
Performance differences between transfer learning and peptide-centric library reconstruction are dependent on software framework. (A) Comparison, relative to the initial library, of precursor identifications and precursor level reproducibility for transfer learning (X) and reconstructed library (circle) across libraries for OpenSWATH (top) and DIA-NN (bottom). Arrows compare the transfer learning and the reconstructed libraries from the same source. The reconstructed library consistently achieves higher reproducibility and, in OpenSWATH, increased precursor identifications. With DIA-NN, initial libraries with a smaller search space identify fewer precursors in the reconstructed library compared to the transfer learning library, likely due to the decrease in sample scope. Differences are more pronounced in OpenSWATH compared with DIA-NN. (B) Comparison of peptide characteristics between the initial (blue), reconstructed (orange) and transfer learning (grey) from OpenSWATH. Characteristics include retention time (top), ion mobility (middle) and relative fragment ion intensity (bottom). Lower variation around 0 indicates higher concordance between the library and sample. The reconstructed library’s RMSD are 2.6-fold lower for RT and 2.1-fold lower for IM compared to transfer learning. This indicates that reconstruction reduces sample-library variation in peptide properties more effectively than transfer learning. RMSD in relative fragment ion intensity remain comparable between approaches.

The higher reproducibility in our library reconstruction workflow can also be explained by its higher peptide property precision. Although both transfer learning and library reconstruction improve peptide property precision, transfer learning does not achieve the same precision across retention time and ion mobility as empirically corrected properties (Figure 3B, Figure S9). Notably, compared to the transfer learning library, the retention time RMSD in the reconstructed library is on average 2.6-fold smaller in OpenSWATH and 1.9-fold smaller in DIA-NN (Figure 3B, Figure S9). As further evidence on the importance of retention time precision, we found that retention time precision explains 42% of the variation in reproducibility across different libraries in OpenSWATH and 57% in DIA-NN (Figure S8). These findings suggest that the higher reproducibility is a result of improvements to both library-sample concordance and peptide property precision.

To separate out the effects of peptide property precision and library-sample concordance, next we analyzed libraries which only filtered peptide precursors and did not correct for peptide properties. Notably, across libraries and software tools, filtering alone always results in higher reproducibility than transfer learning (Figure S10).

When comparing the identifications between the transfer learning and reconstructed workflows, we observed that trends depend on the software and analyte level. This suggests that if maximizing identifications is the primary objective, sample scope and library-sample concordance must be balanced and the optimal balance is dependent on software tool. When using OpenSWATH, peptide-centric library reconstruction always achieved higher precursor and protein identifications with a large effect size relative to DIA-NN (Figure 3A, S11). This demonstrates that with OpenSWATH, maximizing identifications usually requires increasing library-sample concordance and peptide property precision at the cost of sample scope. The strong effect of library-sample concordance on identification rates is likely due to OpenSWATH’s statistical framework. OpenSWATH estimates the proportion of nulls in the library improving statistical power for libraries with strong library-sample concordance. Given that DIA-NN is less sensitive to library quality, it is harder to make generalizable statements comparing identifications between a transfer learning and library reconstruction approach. On the peptide precursor level, the transfer learning approach slightly outperforms library reconstruction in all libraries but the *in silico* library (Figure 3A). This suggests that identification rates in DIA-NN are less dependent on library-sample concordance compared to OpenSWATH, as only the largest library achieves benefits from the reduced library-sample concordance. This is likely because unlike OpenSWATH, DIA-NN does not incorporate estimates on the proportion of nulls into its statistical framework. At the protein level, library reconstruction always identifies more protein groups than transfer learning (Figure S11). These differing trends at the precursor and protein level observed in DIA-NN suggest that the statistical framework at the protein level might be different and may put further weight on the importance of sample concordance and peptide property precision compared to the peptide precursor statistical pipeline.

These trends suggest that the balance between library-sample concordance and sample scope is dependent on software framework and primary objective. In OpenSWATH, library-sample concordance should often be prioritized over sample scope to maximize identifications and reproducibility. Whereas when using DIA-NN, the balance shifts more in favour of preserving sample scope at the cost of library-sample concordance.

### Library Construction Strategies Can Be Adapted to Small Sample Amounts

In smaller sample amounts, the impact of the source of the peptide library on performance increases compared to bulk samples; for example, across a dilution series from 100 ng to 1 ng, as the peptide mass injected decreases, the consistency in the proteins identified across different library sources increases (Figure S12). Notably, the Jaccard index, a similarity metric, between proteins identified across different sample sources is 50% greater in the 100 ng injection compared with the 1 ng injection in both OpenSWATH (0.48 to 0.77) and DIA-NN (0.53 to 0.82) (Figure S12). Given the results above show that our library construction strategies may improve performance in its originating context, we reasoned that this could be extended to alternative cases with a different reference sample. Specifically, we investigated whether a reference sample of larger peptide mass could be used to improve library quality, therefore improving results achievable from an injection with less peptide mass.

In our library construction workflow, we used larger sample amounts (e.g. 25 ng) acquired in DIA as reference samples to construct the library for the smaller sample amounts (e.g. 1 ng). By increasing library quality, peptide-centric library reconstruction enables peak groups of low intensity to be confidently identified (Figure 4A).

**Figure 4:**
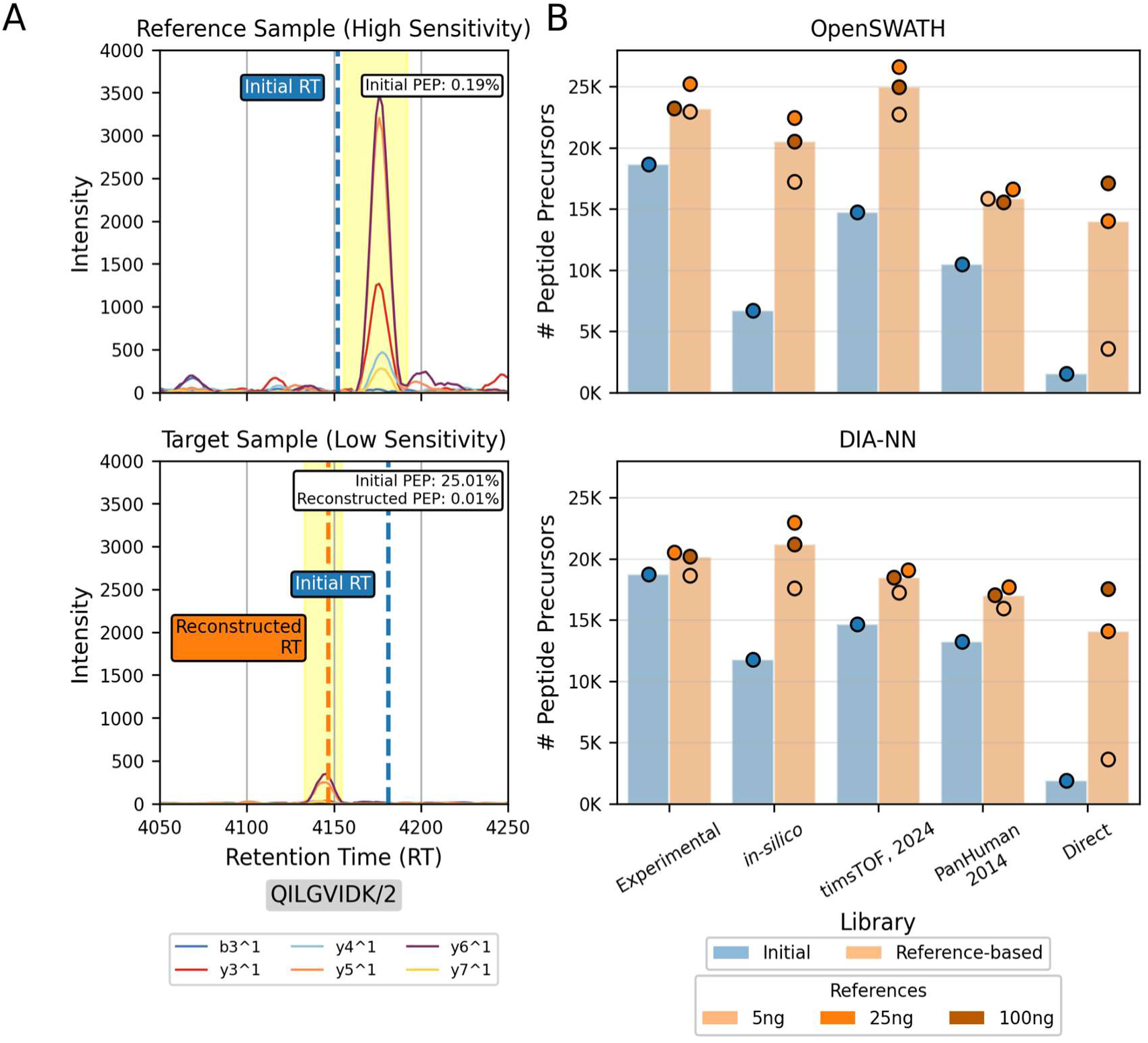
Reference-based library construction using larger sample amounts increases precursor identifications across library sources and software tools. (A) Example of a peptide precursor, *QILGVIDK^2+^,* that is only confidently detected after peptide-centric library reconstruction. Searching for this precursor with the initial *in silico* library only yields a confident identification in the reference 25 ng sample (PEP = 0.19%) and not in the 1 ng sample (PEP = 25.01%). However, the library reconstructed with the 25 ng reference enables a confident identification (PEP = 0.01%) in the target 1 ng dilution. Library retention time accuracy improves after reconstruction. Note that although the initial libraries are the same, the retention times are different due to internal nonlinear retention time calibration before analysis. (B) Number of peptide precursors identified at 1 ng sample injection across libraries for OpenSWATH (top) and DIA-NN (bottom) comparing the initial (blue) and reference-based (orange) versions. Points represent the number of peptide precursors obtained using different reference samples for library construction, ranging from 5 ng to 100 ng, with the bar’s height representing the median identifications across the different reference dilutions. Applying the spectrum-centric library construction workflow on the direct library achieves over a 9-fold increase in precursor identifications across both DIA-NN and OpenSWATH. The optimal reference for maximizing peptide precursor identifications is 25 ng for peptide-centric reconstruction and 100 ng for spectrum-centric construction.

Remarkably, increasing library quality using a reference-sample mitigates the discrepancy in protein identifications across different library sources, achieving a Jaccard index similar to what was achieved by the 100ng dilution without our reference-based library construction (Figure S12). Improvements in Jaccard index were most pronounced in the smallest sample injection amounts with the 1 ng injection exhibiting a 45% improvement in Jaccard index after increasing library quality for both OpenSWATH (0.48 to 0.70) and DIA-NN (0.53 to 0.77). This suggests that incorporating reference samples into library construction can compensate for the challenges of selecting the best library source when dealing with small sample amounts.

In other words, the effects of reference-based construction on proteome coverage were most pronounced in the smallest sample amounts (Figure S13). At the 1 ng dilution, we observed that, in contrast to bulk samples, incorporating a reference sample improves identifications across all libraries and software tools with a median increase of 60% at the peptide precursor level and 40% at the protein level (Figure 4B, Figure S14). This consistent increase in identifications suggests that a high quality library is important at low sample loads regardless of the software framework used. Effects were most pronounced in the lowest quality library sources and lowest injection amount, which in this case is the direct library at the 1 ng dilution. In this workflow, DIA-reference spectrum-centric library construction increased peptide precursor identifications over 9-fold, such that the number of identifications of the new library were similar in magnitude to the peptide-centric approaches (Figure S13).

One of the major challenges with this approach is determining the optimal reference sample load. Although larger sample amounts offer the potential for greater sample scope as they lead to larger libraries, smaller sample amounts result in a library with higher library-sample concordance. To test this, we performed library construction with three different reference samples: 5 ng, 25 ng and 100 ng. For the peptide-centric library reconstruction approach, reconstruction with the 25 ng reference resulted in the deepest coverage (Figure 4B, Figure S14). Considering that the peptide property precision remains relatively consistent across the different reference samples (Figure S15), this suggests that sample scope and library-sample concordance primarily explain the difference in performance between the reference sample loads. Therefore, the 25 ng reference likely leads to the best results because the 5 ng injection does not provide sufficient material for high quality reconstruction while the library reconstructed using the 100 ng injection has lower concordance with the target sample. For the spectrum-centric library construction approach, the 100 ng injection was the most suitable reference (Figure 4B, Figure S14) indicating that the lower injections do not provide sufficient signal to directly construct a library. Overall, this suggests that it is important to select the correct reference for library construction and that the best reference depends on the library construction strategy.

### Library Construction in Single-Cell-Equivalent Protein Quantities

Given that our reference-based library construction strategy excels at limited sample amounts, we next aimed to demonstrate the potential for this workflow to be adapted to single cell proteomics. To test this, we used the timsTOF-SCP, which can achieve more than 10-fold increase in sensitivity compared to the instrument used above.^6^

This dilution series contained six steps ranging from 5000 pg to 100 pg, with nine technical replicates per dilution step. We used injections from a larger sample load as reference to construct the library for smaller sample amounts. Importantly, this dataset did not contain an associated offline fractionated experimental library simulating conditions where sample amounts are too limited to acquire one.

First, we examined the effects of improving library quality across the dilution steps. For simplicity, we initially used the 5000 pg replicate with the highest proteome coverage as the reference as this dilution step can serve as a reference for all other dilutions. Similar to above, we found that incorporating the reference sample into library construction had the greatest effect on precursor identifications at the smallest sample amounts (Figure S16). Furthermore, we also found that increasing library quality increases the similarity in proteins identified across library sources. This effect was most pronounced at the smallest sample amounts (Figure S17). For example, when analyzing with the initial libraries using DIA-NN, the 100 pg dilution contains many protein identifications unique to a specific library source with Jaccard indices across libraries of 0.41. After incorporating the reference sample into library construction, the Jaccard index improved 75% to 0.72 which is greater than what was achieved in the initial library with the 5000 pg sample injection (Jaccard index 0.71). This demonstrates that reference-based library construction can decrease variation in results attributed to different library sources effectively mitigating the effects of a lower quality library source.

Considering the effects were most beneficial at the smallest sample amount of 100 pg, next we focused our attention on optimizing this workflow. First, we systematically tested our library construction strategy against all possible reference samples using the replicate with the most comprehensive coverage at each dilution step to perform library construction. Interestingly, this unveiled that the trade-off between reproducibility and proteome coverage is dependent on the reference sample load. More specifically, higher load reference samples prioritize proteome coverage over reproducibility (Figure S18). For example, consider the analysis of a 100 pg sample with the *in silico* library and OpenSWATH: A library reconstructed with the 250 pg as a reference increases precursor identifications by 78% and Jaccard index by 58%. In contrast, library reconstruction with the 5000 pg increases precursor identifications 3.5-fold, but the average Jaccard index is practically unchanged (percent increase <2%). These trends are likely observed because larger sample amounts prioritize sample scope over library-sample concordance, enabling deeper proteome coverage at the cost of reproducibility. This is consistent with previously observed trends, which find that libraries more closely related to the target sample, favour reproducibility over comprehensiveness.^48,49^ In this context, we found that the 500 pg dilution provided a good trade-off between reproducibility and proteome coverage across all library sources tested (Figure S18). Furthermore, we did not observe major differences in the peptide property precision across different reference samples used (Figure S19).

When using a 500 pg sample as a reference for the 100 pg results (Figure 5A), we observed that in this low input application, incorporating the 500 pg reference into library construction leads to a consistent improvement in reproducibility and proteome coverage across libraries and software tools (Figure 5C, Figure S20). These results reinforce that, unlike at bulk sample loads, both software frameworks benefit from improved library quality in low-input settings. Furthermore, looking at a specific chromatogram, it is evident that increasing library quality using the 500 pg reference enables the confident detection of peptides with low signal (Figure 5B). In a spectrum-centric context, peptide precursor identifications increased 6-fold (∼1000 to ∼6000). Notably, using the reconstructed timsTOF library with OpenSWATH, we identified over 15,000 peptide precursors (2480 protein groups) from a 100 pg sample load, a 90% increase compared to before library reconstruction (8024 peptide precursors). Interestingly, this improvement is independent of peptide modifications, as unmodified peptides exhibit a similar proportional gain from 6187 to 11,653 stripped peptides (88% increase). We also found that across all libraries and software tools, library construction consistently increased the number of quantitatively precise peptide precursors quantified across all nine replicates (Figure 5D, Figure S21). Notably, using OpenSWATH, *in silico* library reconstruction yields a 1.8-fold increase in the number of commonly identified peptide precursors quantified with a CV in intensity of less than 20% (815 to 1480).

**Figure 5:**
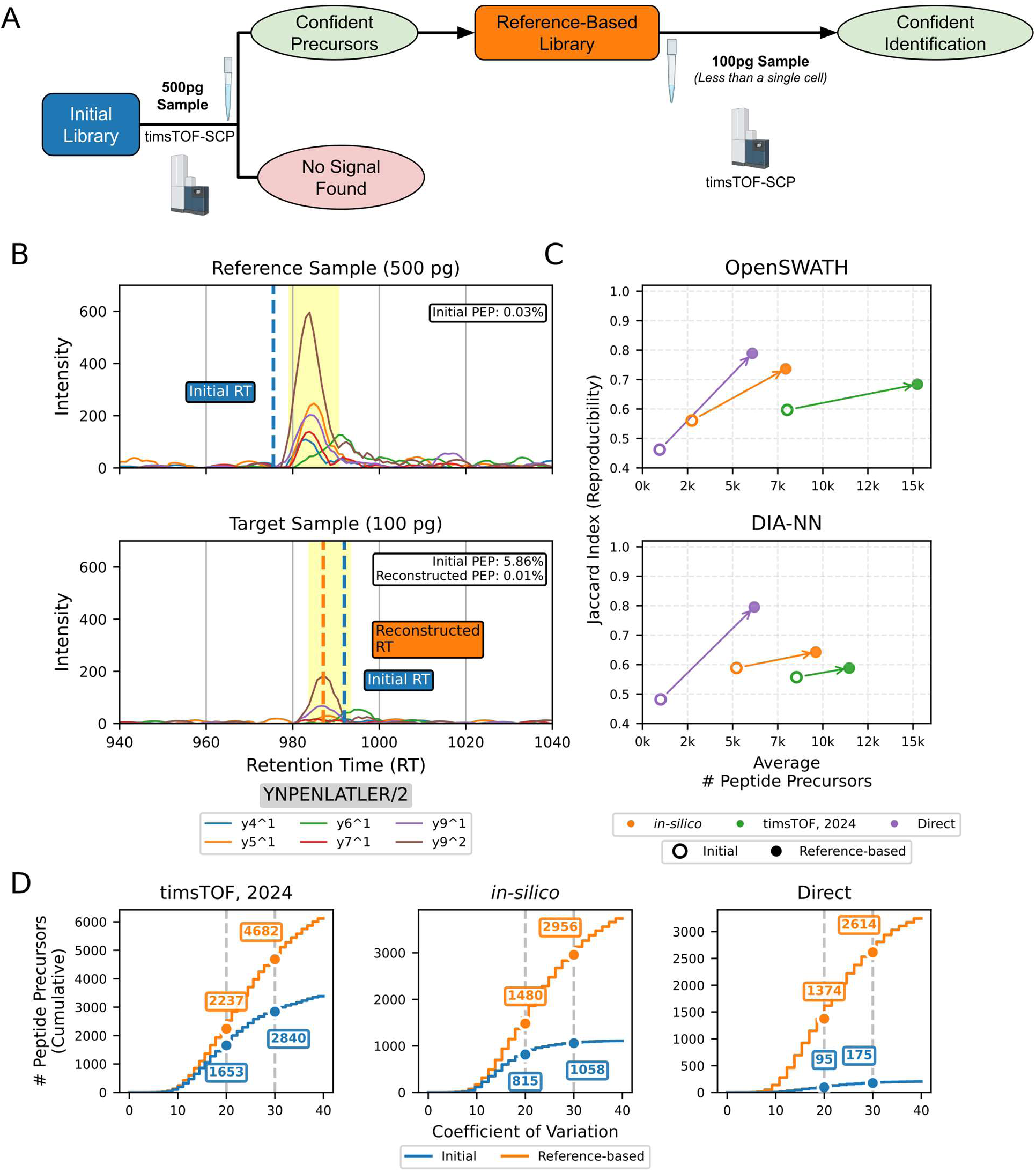
Library construction improves precursor identifications and reproducibility in single-cell-equivalent HeLa cell digests across both software frameworks. (A) Schematic illustrating the library construction workflow with a 100 pg target sample. A 500 pg sample is analyzed with the initial library to create a new library. This new library is then used to analyze the results from a 100 pg protein sample. Created with BioRender.com (2026). (B) Example of a peptide precursor, *YNPENLATLER^2+^*that is only confidently detected after library reconstruction. This precursor is confidently identified with the initial *in silico* library in the 500 pg sample (PEP = 0.03%) however not with the 100 pg sample (PEP = 5.86%). Using the reconstructed library enables a confident identification in the 100 pg sample (PEP = 0.01%). Note that although the initial libraries are the same, the retention times are different due to internal nonlinear retention time calibration. (C) Comparison of precursor identifications and reproducibility averaged across the 9 replicates. Reference-based library construction consistently increases the number of precursors identified and reproducibility across all software tools with effects most pronounced in the spectrum-centric approach. Two replicates analyzed with the initial direct library were excluded from mean calculations since no peptide precursors passed the 1% protein FDR threshold. Notably, reference-based library construction using a spectrum-centric approach leads to a 6-fold improvement in peptide precursor identifications over constructing the library directly from the sample. Furthermore, with OpenSWATH, over 15,000 precursor identifications can be achieved after library reconstruction of the timsTOF, 2024 library. (D) Cumulative distribution of commonly identified peptide precursors (1% FDR) binned by their CV in intensity from OpenSWATH results. The reference-based libraries quantify on average 850 and 2000 more peptides with an intensity CV less than 20% and 30% respectively across all library sources.

Overall, this demonstrates that our referenced-based library construction workflow increases library quality leading to improved proteome coverage, reproducibility and quantification precision in low-input experiments.

## Discussion

In single cell and low-input proteomics, one of the greatest difficulties when using DIA is generating a library of sufficient quality. This challenge leads to a plethora of ad hoc approaches to library generation in single cell and low input settings, with vastly different qualities of the resulting libraries.^6,19–22,50^ Here, we implement a systematic approach to evaluate different strategies for library construction and show that the choice of library affects performance, especially in small sample amounts. We provide a library construction strategy and show that incorporating our strategy minimizes the impact of the initial library, particularly benefiting low input proteomics applications. Briefly, this workflow involves a preliminary analysis with the reference sample to construct a library, then using this library to analyze the sample of interest. This strategy aims to maximize quality at a minor cost in sample conservation because additional injections are required for library construction. Reference samples are selected to be closely related to the target sample but of higher quality resulting in data with greater signal or higher selectivity. We show that this strategy can be employed in both peptide and spectrum-centric approaches, the former of which can be conducted in a completely open-source framework. We demonstrate that our strategy can be applied across a variety of library sources of varying quality and across two different software frameworks, DIA-NN and OpenSWATH. During evaluation, we purposely did not use any MBR approaches to ensure that these strategies could be generalized to applications which contained more heterogeneous samples, such as single cell proteomics, where MBR might not be appropriate.^26^

First, we applied this strategy in its originally developed context where the reference sample consists of GPF samples.^28,29^ The higher selectivity of the GPF reference improves library quality by reducing the search space and ensuring peptide properties are more consistent with the target sample. This results in the target sample operating as if it too had an increase in selectivity; effectively transferring the increased selectivity of the reference sample to the target sample. Here, we demonstrate that both OpenSWATH and DIA-NN exhibit improved reproducibility with the reference-based library compared to the initial library. This improvement can likely be attributed to the improved library-sample concordance and peptide property precision. This is consistent with the observation that reference-based library construction achieves higher reproducibility than a transfer learning approach, which does not improve concordance and offers fewer gains in precision. However, the software tools differ in their effect size and trends in identification counts. OpenSWATH shows improved identification counts regardless of the initial library, a consistent trend with prior studies.^28^ In contrast, DIA-NN only shows pronounced increases in identifications for libraries with large search spaces. One explanation for these differing trends is the different statistical approaches employed by these tools. On one hand, OpenSWATH estimates the proportion of nulls in the library when computing q-value;^42,51^ and because reference-based libraries contain a low proportion of true negatives, this approach leverages that increased statistical power into a substantial increase in proteome coverage. In contrast, DIA-NN is described as not estimating the proportion of nulls in the library when computing q-values,^43^ meaning that it cannot take full advantage of very high-quality libraries. This highlights a key difference between the software tool design: OpenSWATH is more sensitive to library quality compared to DIA-NN. On one hand, this means only DIA-NN can achieve similar results from both a high quality experimental library and a low quality *in silico* library. However, the benefits of ultra high-quality libraries, generated with reference samples, can only be exploited fully by OpenSWATH. Our library reconstruction workflow notably closes the performance gap by elevating OpenSWATH’s performance with *in silico* libraries to at the level of DIA-NN. We hypothesize this behaviour reflects differences in their design principles. OpenSWATH was designed to analyze DIA data using a targeted proteomics framework, where peak groups are chosen using fixed criteria and machine learning is used afterwards for statistical error rate estimation.^8^ In contrast, according to the literature, DIA-NN iteratively performs peak group selection based on target-decoy competition,^43^ allowing for peak selection to evolve in a data driven manner. This may explain why DIA-NN appears less sensitive to library quality in statistical error rate estimation and its lack of ability to exploit a high quality library.

An alternative approach to conduct peptide-centric library reconstruction is using a two pass approach, where the same dataset is used to reconstruct the library and for subsequent analysis. However, this violates the assumption that decoys are drawn from the same distribution as false target signals because targets are pre-selected for high scores in the first step.^52–55^ In contrast, our approach reconstructs the library using an independent dataset, thereby avoiding this specific violation.

However, since our FDR framework assumes all entries in the library are true, our approach may transfer false positives in the reference-based library.^28,56^ In practice, this can be controlled by adjusting the reference sample FDR cutoff to be included in the new library. Although this is also a concern for libraries constructed from using DDA samples, this problem is pronounced when constructing the library from reference samples more closely related to the target such as GPF or a larger sample load. This is because in higher quality libraries constructed using references more closely related to the target sample, the majority of false signals present are highly similar to true signals, making them difficult to distinguish. This can consequently cause FDR estimation to be more liberal.

To address this, we ensured that all entries of the reference-based constructed library were identified at 1% precursor, peptide (if applicable) and protein-level FDR in the reference sample. Precursors were also removed if they contained fewer than 4 transitions. However, as we observed with DIA-NN (Figure 3A), stringently filtering the library limits the scope of the library which may limit the proteome coverage achievable. Therefore, a new statistical framework that accounts for false entries in the library, such as the one proposed by Frejno et al., 2025 ^52^ may increase the impact of reference-based library construction on proteome coverage.

It is important to note that even when statistical frameworks are working as expected, using strong priors may increase the number of noisy signals which are confidently identified.^57^ In this context, this means that a high-quality library allows detection of peptides close to the limit of detection which a lower quality library does not. This does however highlight limitations with the current statistical approaches and suggests that new approaches may be required to further distinguish between peptides worth examining in follow up studies.^58,59^ Considering this, it is important to also evaluate the impact of reference-based library construction on quantitative precision. We demonstrate that peptide-centric library reconstruction can change the peak group being chosen leading to more precise quantification. Overall, we observe that our library construction workflow consistently improves the number of quantitatively precise peptide precursors across different applications, libraries and software tools.

Transfer learning has shown great promise in a large number of applications including proteomics.^38,60–63^ Instead of using the GPF runs to directly reconstruct the library, we evaluated using a transfer learning approach to refine *in silico* models for predicting peptide properties. This approach has two major implications compared to our peptide-centric library reconstruction approach. First, in transfer learning, peptide properties are refined using a model rather than empirical evidence directly, meaning that peptide property precision is reduced. It is important to note that if transfer learning uses the target dataset the model can outperform empirical evidence;^60^ however, this involves conducting transfer learning on the same dataset used for analysis which may violate statistical assumptions as discussed above. Secondly, the search space is not reduced. Although this enables the detection of peptides that were not confidently detectable in the GPF runs, reducing the search space is important to improve reproducibility for all software frameworks and to increase proteome coverage with OpenSWATH. Trends observed with OpenSWATH are consistent with previous findings on the importance of filtering the library for increasing proteome coverage.^60,64^ Considering DIA-NN is less sensitive to library-sample concordance, the trade-offs between the transfer learning approach and the library reconstruction strategy is dependent on the library source. At the peptide precursor level, if the initial library already has high library-sample concordance, DIA-NN achieves higher identifications with transfer learning compared to library reconstruction as transfer learning does not affect sample scope. Interestingly, the library reconstruction approach slightly outperforms DIA-NN in the number of proteins identified suggesting that increasing library-sample concordance is important for improving confidence on the protein level.

Although improved instrumentation may enable acceptable results across a wide range of libraries of various qualities, especially with software tools less sensitive to library quality, the effect of library quality becomes evident when pushing the instrumentation to its limit of detection. For instance, lower sample amounts lead to increased variability in the proteins detectable across different library sources. Fortunately, our library construction workflow can be easily adapted to low-input experiments by using higher input injections as the reference sample to construct the library. By increasing library quality, the peak groups in the target sample are scored as if they had the sensitivity of the reference sample, effectively transferring the sensitivity from the reference sample onto the target sample.

In this approach, only a single extra injection is required to vastly improve the quality of the library, improving precursor identification rates in both DIA-NN and OpenSWATH to be comparable to an offline fractionated experimental library. Therefore, our library construction strategy is a viable approach for achieving the coverage of an offline fractionated library in applications where it is not feasible to construct an offline fractionated library.

Finally, we applied our strategy to single cell equivalent proteome injections of just 100 pg of peptide mass. In this approach, a reference sample of 500 pg was used to construct a high-quality library for low input samples. Using this reference sample ensures that the reference sample is closely related to the target sample, which is important to consider when dealing with small sample amounts.^65^ The effects of library construction were most pronounced in our spectrum-centric strategy. In sub-single cell equivalent injections (100 pg), we observed a 6-fold improvement across both OpenSWATH and DIA-NN. Similar to bulk sample amounts, in our peptide-centric workflow, the most pronounced improvements were observed when reconstructing the *in silico* library. However, unlike in bulk samples amounts, reference-based library construction consistently increased identifications across both software tools, yielding a 34-190% increase in peptide precursors for all libraries tested. This highlights that despite DIA-NN being relatively robust against the effect of library quality in bulk sample amounts, in small sample amounts high quality libraries are a necessity. Furthermore, in our fully open source peptide-centric pipeline using OpenSWATH, we detected over 15,000 peptide precursors (2480 protein groups) from 100 pg of sample. Overall, considering the low cost in sample conservation and large improvements observed across all workflows in small sample amounts, we highly recommend using a library construction approach in single cell proteomics and other applications where sample amounts are limited.

## Conclusion

We conclude that our reference-based library construction strategy is an effective strategy at improving proteome coverage, reproducibility and quantitative precision in both bulk and sample limited applications. In bulk sample amounts, the improvements observed are more pronounced in OpenSWATH, however in small sample amounts notable improvements are observed in both DIA-NN and OpenSWATH. By increasing library quality, the peak groups are scored as if they had higher sensitivity or selectivity effectively transferring the sensitivity or selectivity from the reference sample onto the target sample. Our approach is adaptable to different libraries, software tools and analyses.

In bulk samples, high selectivity GPF experiments can serve as a reference to maximize selectivity achievable in the target sample. In this context, the improvements observed are most pronounced in the OpenSWATH framework whose strong dependency on library quality provides a controlled and interpretable framework for assessing the effect of the library used. Using OpenSWATH, our strategy clearly outperforms transfer learning approaches through its combination of reducing the search space to only high confident peptides, ensuring highly precise empirically derived peptide properties. Although future work is needed for further statistical validation, our approach avoids limitations of other two pass approaches by using an independent dataset and stringent filtering criteria when constructing the library.

The benefits of our library construction approach are most pronounced across both software tools when sample amounts are limited. In single cell equivalent (100 pg) injections, our strategy identifies over 15,000 peptide precursors with OpenSWATH. This strategy could naturally extend to single cell proteomics by using a multi-cell protein digest as a reference, where we anticipate it providing substantial benefits.

Overall, these findings highlight the need for using high quality libraries, specifically in software frameworks sensitive to library quality and when sample amounts are limited. Here, we present a solution for improving library quality using high quality reference samples. These high-quality libraries can be used to maximize the information extractable from a single mass spectrometry injection improving proteome coverage, reproducibility and quantification accuracy.

## Data Availability

The proteomics datasets are available via MassIVE data repository with the identifier MSV000102409.

## Acknowledgements

J.C. acknowledges financial support from the Ontario Graduate Scholarship. J.C., M.G., and H.L.R were funded by the Canadian Institutes of Health Research (CIHR) (#506078). H.L.R. is the Canada Research Chair in Mass Spectrometry-based Personalized Medicine. A.C.G. was funded by the Terry Fox New Frontiers Program Project Grant (PPG). Mass spectrometry data was acquired at the Network Biology Collaborative Centre Proteomics Facility (RRID: SCR_025375) at the Lunenfeld-Tanenbaum Research Institute, Sinai Health Research. The facility is supported by the Canada Foundation for Innovation and the Ontario Ministry of Colleges, Universities, Research Excellence and Security (MCURES).

