## Supplemental Figures for "Reference-Based Library Construction Improves Performance in Low-input Workflows"

### Supporting Information - Refined Library Searching Improves Performance in diaPASEF Workflows

**1 Donnelly Centre for Cellular and Biomolecular Research, University of Toronto, Toronto, Ontario, Canada**

**2. Department of Molecular Genetics, University of Toronto, Toronto, Ontario, Canada**

**3. Lunenfeld-Tanenbaum Research Institute, Toronto, Ontario, Canada**

**4. Department of Computer Science, University of Toronto, Toronto, Canada**

*** Corresponding author**

**Corresponding**

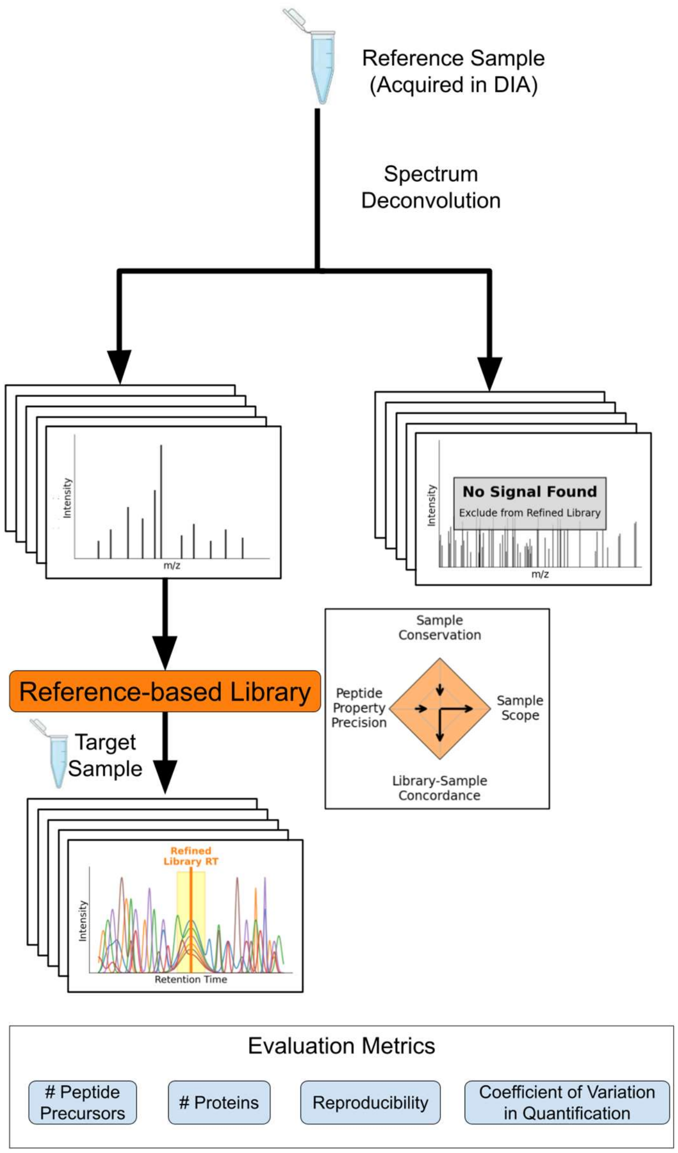


**Figure S1: Overview of the spectrum-centric library construction workflow.** In this workflow, the library is constructed directly from a DIA reference sample in a spectrum-centric manner. This includes spectrum deconvolution and a database search. The reference-based library is used with the target sample to search in a peptide-centric manner. In this approach, the reference-based library would have deeper coverage and higher similarity with the sample than if the library was constructed directly from the target sample. Created with BioRender.com (2026).

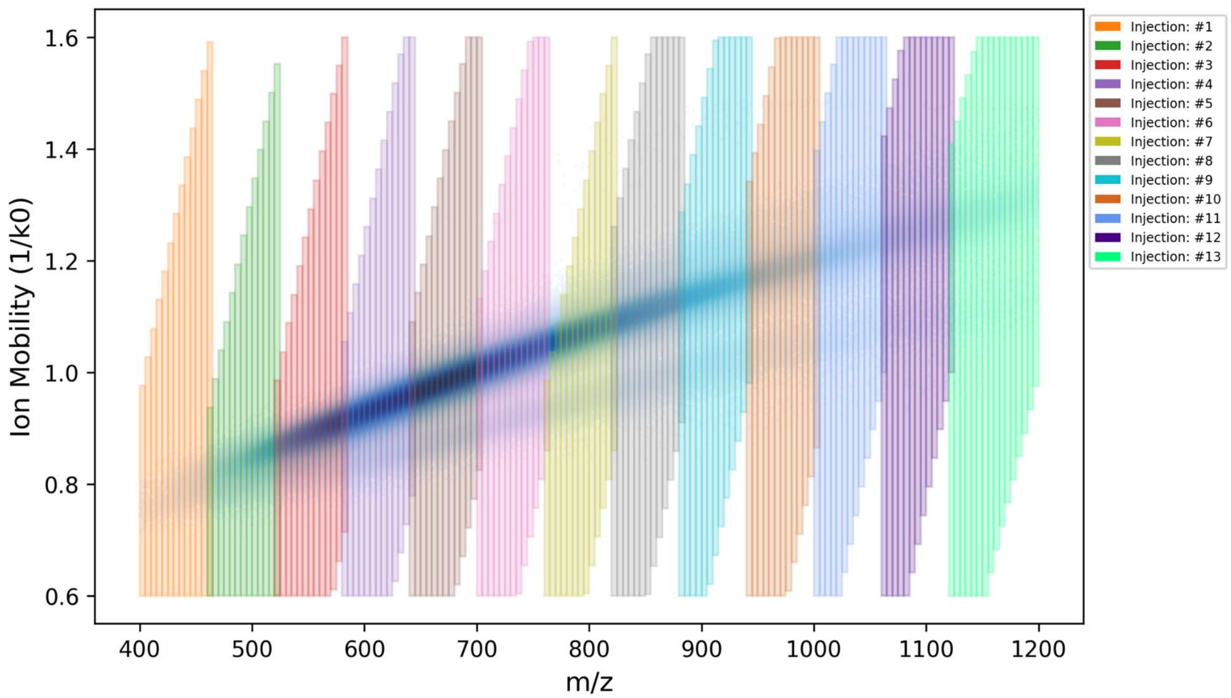


**Figure S2: Overview of the gas-phase fractionation (GPF) sampling scheme.** The GPF sampling scheme used in this experiment overlaid on top of a distribution of the peptide precursors in the experimental library. The first 11 injections all cover a 60 m/z region in 5 m/z isolation windows. The last 2 injections cover an 80 m/z region with 5 m/z isolation windows.

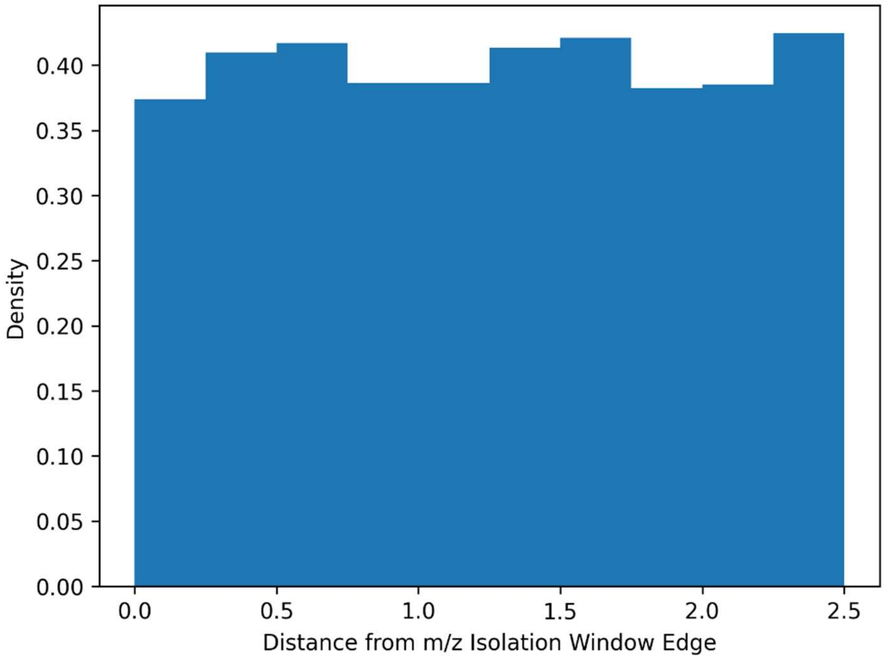


**Figure S3: m/z isolation window edge effects are not present in the GPF sampling scheme.** Distribution of identified peptide precursors by their distance to the edge of the window. There is no drop in identifications at the edges, indicating that edge-related detection loss is not present.

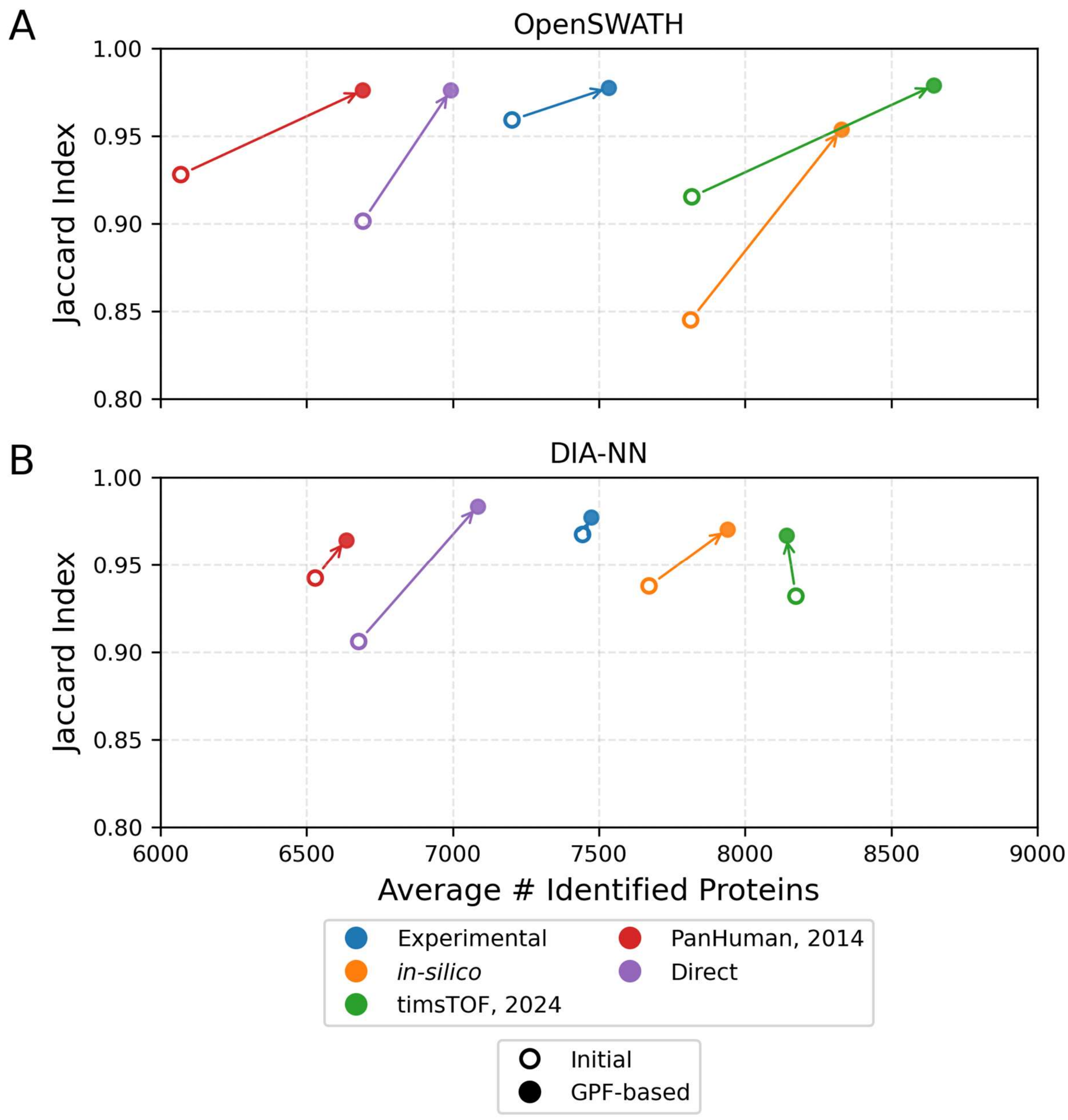


**Figure S4: GPF-based library construction usually improves protein group identifications and reproducibility across libraries and software tools.** Average number of identified protein groups across three replicates against overall reproducibility (Jaccard index) across libraries for two software tools: (A) OpenSWATH and (B) DIA-NN. The results from the initial library are represented by a hollow circle, and the results from the GPF-based library are colored with a filled-in circle. Different colors represent the different libraries tested. Arrows compare the relationship between the initial and GPF-based library for each library source. In general, arrows point to the upper right indicating an increase in protein identification and reproducibility. OpenSWATH exhibits a 3.3-fold greater improvement in identifications and 1.8-fold greater improvement in reproducibility.

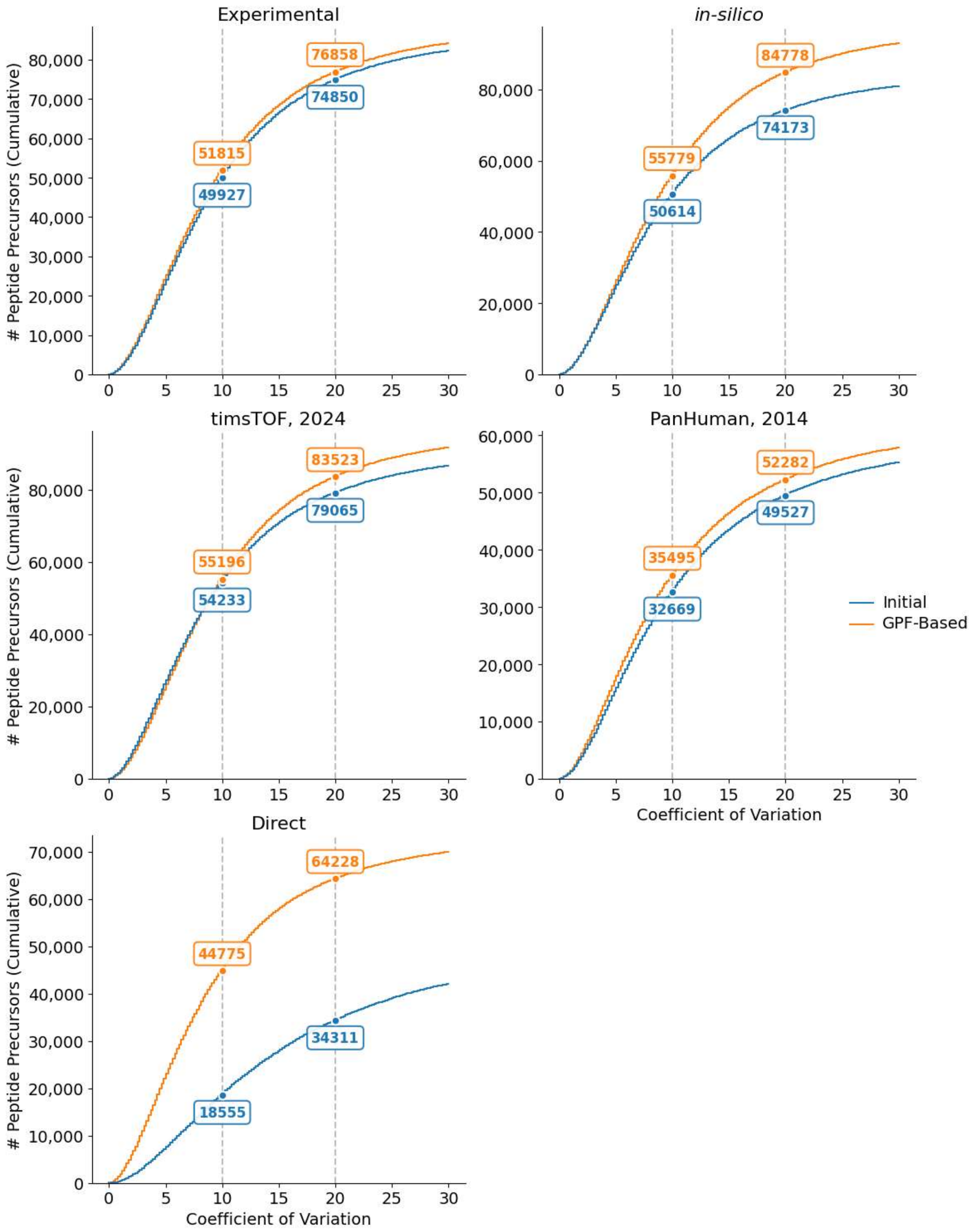


**Figure S5: GPF-based library construction improves the number of quantitatively precise peptides in DIA-NN.** Cumulative distribution of precursors consistently identified in all three replicates at 1% FDR from DIA-NN ranked by their coefficient of variation (CV) in precursor intensity for the initial (blue) and GPF-based (orange) libraries. The GPF-based libraries quantify on average 7.5K and 10K more peptides with an intensity CV less than 10% and 20% respectively across all library sources.

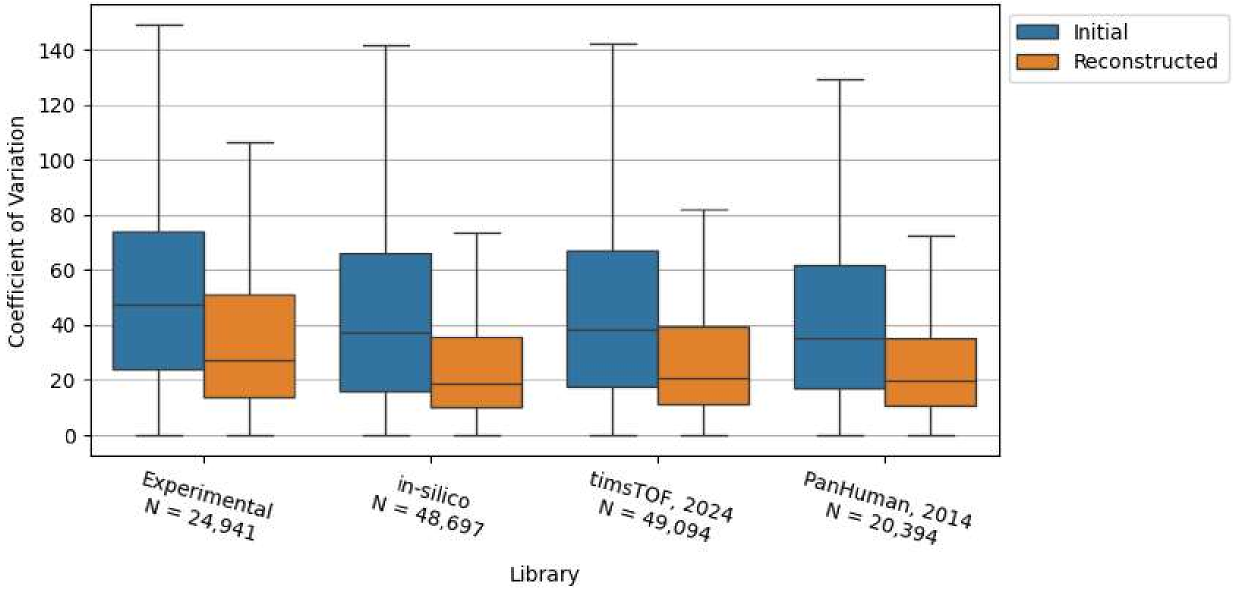


**Figure S6: Newly identified peptide precursors after peptide-centric library reconstruction have lower CV compared to features found in the initial library.** Comparison of quantitative precision of peptide precursors only commonly identified at 1% FDR in all triplicates after GPF library reconstruction. The CV of the reconstructed peptide precursors are shown in orange and initial peptide precursors (which may not necessarily be confidently identified) in blue. These peptide precursors exhibit a lower CV in intensity after library reconstruction. This indicates that library reconstruction causes the search engine to locate the correct peak boundaries which could not be located with the initial library. Only OpenSWATH results are shown since the DIA-NN output does not output a feature for all library entries.

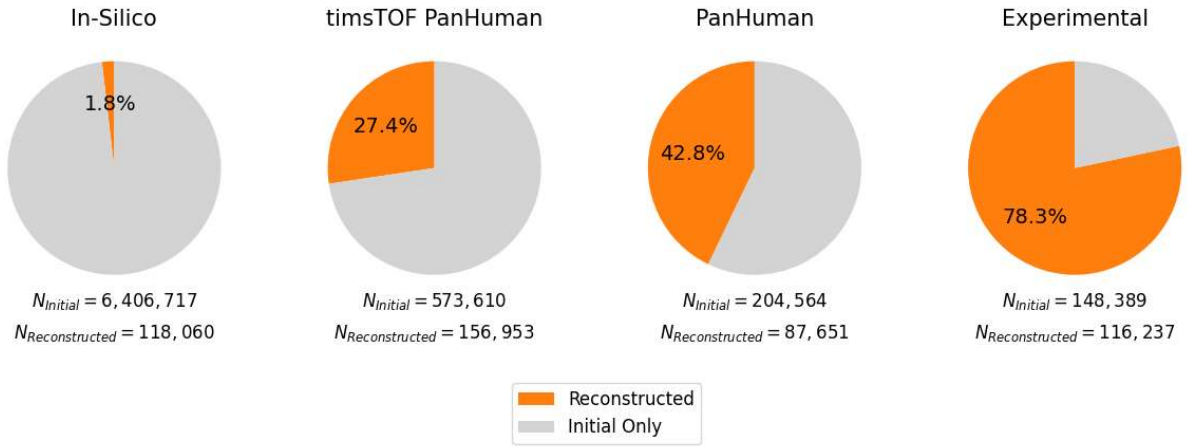


**Figure S7: Peptide-centric library reconstruction reduces search space.** Pie charts illustrating the proportion of peptide precursors in the initial library which are contained in the reconstructed library across different library sources. Larger libraries, which are less concordant with the target sample, have a higher degree of filtering compared to libraries with less peptide precursors.

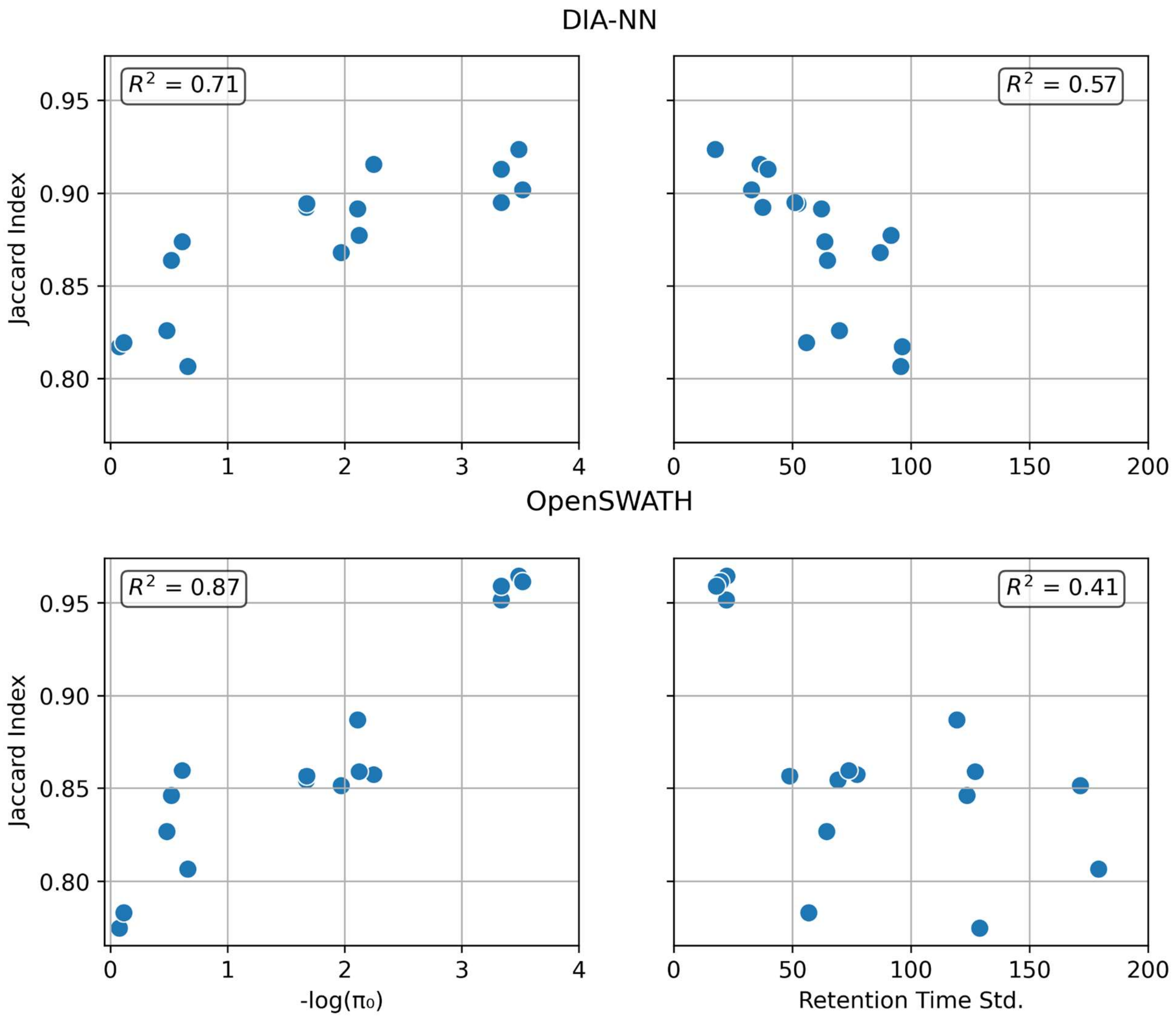


**Figure S8: Variation in reproducibility can be largely explained by estimated proportion of null hypothesis** ($\pi_0$) **and retention time precision.** Scatterplot between Jaccard index and the proportion of true positives metric, measured as -log($\pi_0$) for DIA-NN (top) and OpenSWATH (bottom). $\pi_0$ is defined as the proportion of library precursors that are not detectable in the target sample and is estimated using the Storey-Tibshirani approach in Pyprophet. Points represent different libraries and different refinement strategies including initial, reconstruction and only filtering. In both software tools, the variation in reproducibility can be largely explained by the proportion of true positives in the library. This indicates why transfer learning, which does not filter the library thus not altering the proportion of true positives in the library consistently achieves lower reproducibility compared to the reconstructed library.

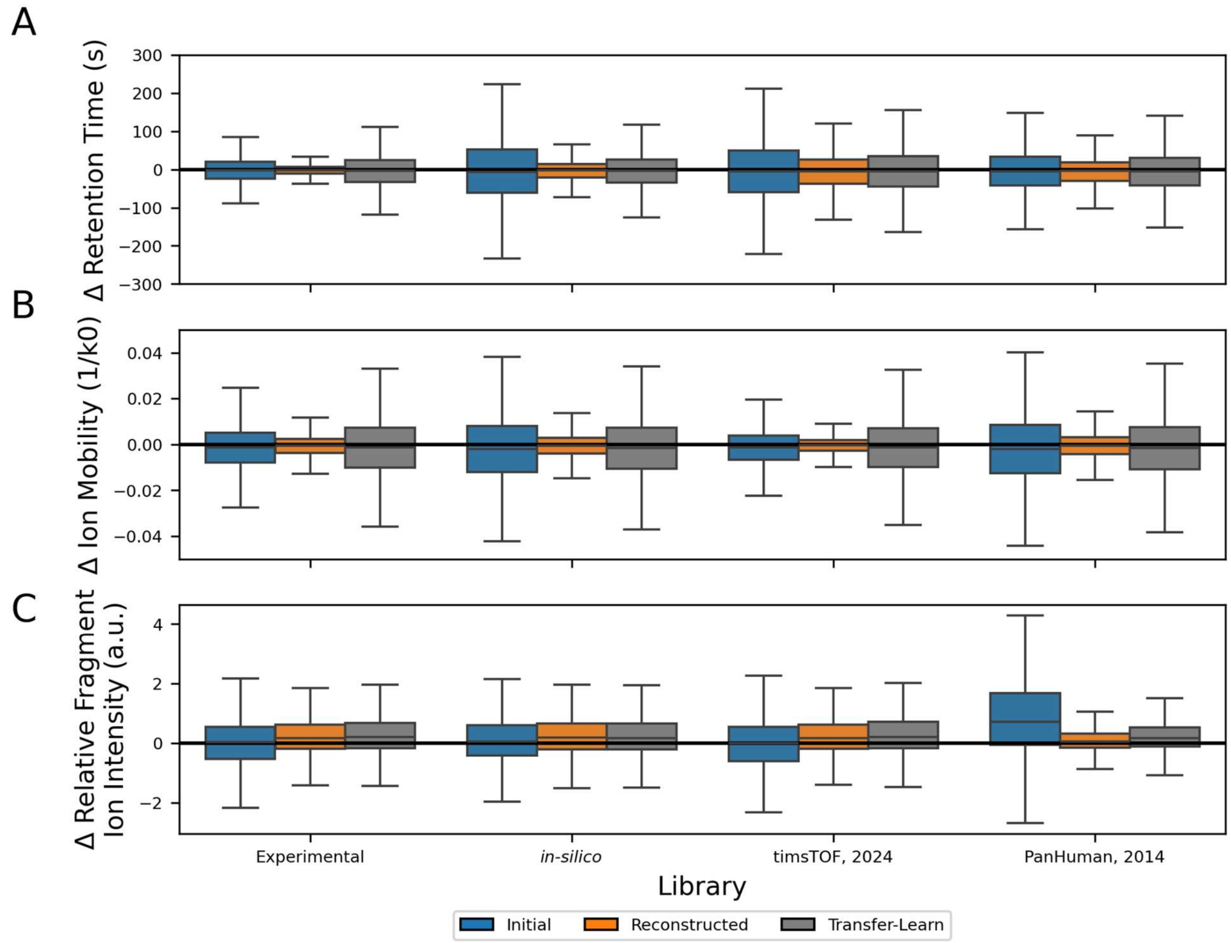


**Figure S9: Comparison of peptide properties across different library refinement strategies analyzed with DIA-NN.** Boxplot showing the experiment-library variation across (A) retention time (B) ion mobility and (C) relative fragment ion intensity across the initial (blue), reconstructed (orange), and transfer-learn (grey) libraries analyzed with DIA-NN. Higher library-sample agreement is represented by boxplots with a low variation centered around zero. Across all libraries and peptide characteristics reconstruction decreases the variation between the experiment and library. Library reconstruction always achieves lower variation compared to the initial library. The reconstructed library's RMSD are 1.9-fold lower for RT and 2.6-fold lower for IM compared to transfer learning. RMSD in relative fragment ion intensity remain comparable between approaches.

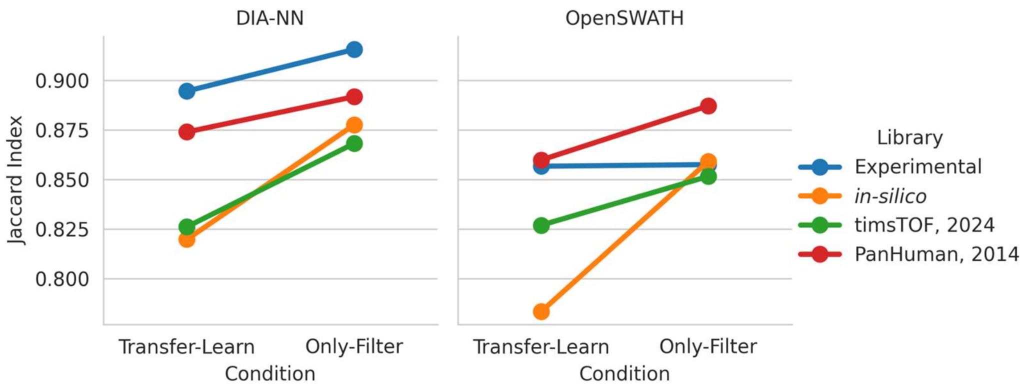


**Figure S10: Filtering alone results in higher reproducibility compared with transfer learning.** Jaccard index for the transfer learning and only filtering library refinement strategies across different libraries for DIA-NN (left) and OpenSWATH (right). Across all initial libraries, only filtering the library leads to higher reproducibility than a transfer learning workflow. Effects are most pronounced in larger libraries.

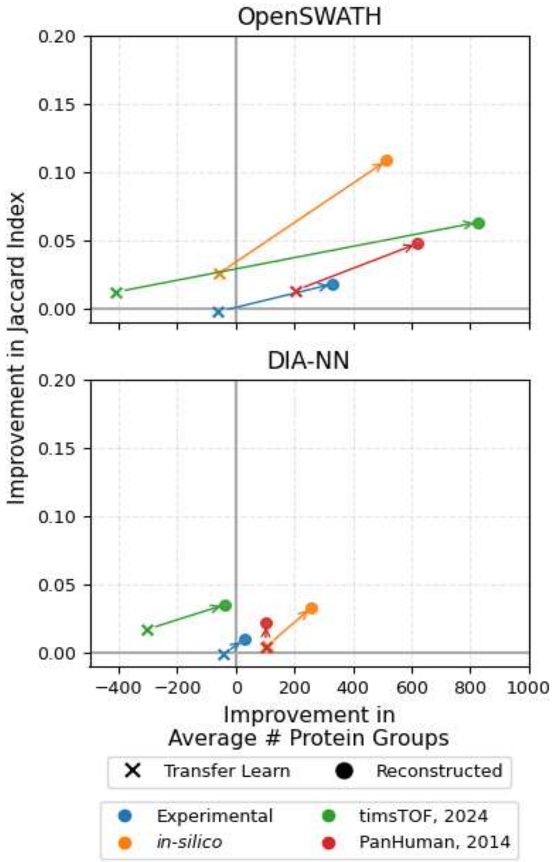


**Figure S11: At the protein level, library reconstruction achieves higher identifications and reproducibility than transfer learning.** (A) Comparison, relative to the initial library, of protein group identifications and protein group level reproducibility for transfer learning (X) and reconstructed library (circle) across libraries for OpenSWATH (top) and DIA-NN (bottom). Arrows compare the transfer learning and the reconstructed libraries from the same source. Differences in reproducibility and identifications are 2.5-fold and 5.3-fold more pronounced in OpenSWATH compared to DIA-NN.

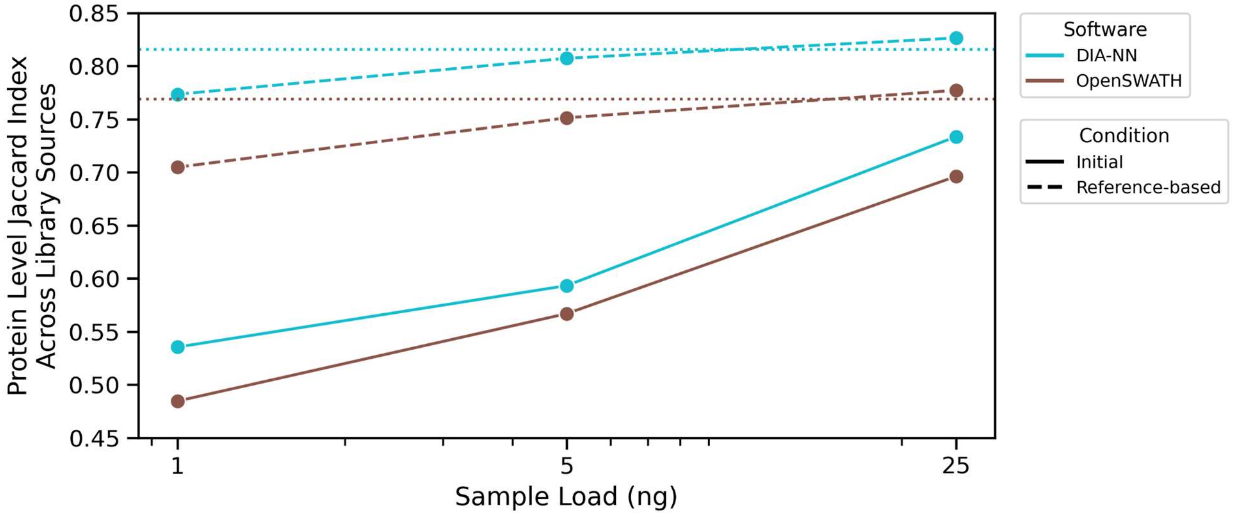


**Figure S12: Library construction from a 100 ng reference sample decreases protein-level variation between libraries.** Comparison of proteins identified for the initial (solid) and reference-based (dashed) libraries for DIA-NN (teal) and OpenSWATH (brown) across the dilution series. x-axis is logarithmically scaled. Dotted lines indicate the reproducibility achieved at the 100 ng sample injection. Comparisons are done at the protein-level to account for the different search spaces in different libraries. Across the initial libraries, reproducibility across different library sources is largely dependent on the sample load with the smallest sample amounts achieving the lowest reproducibility. Reference-based library construction largely mitigates this dependency enabling a relatively consistent set of protein groups across library sources regardless of sample load.

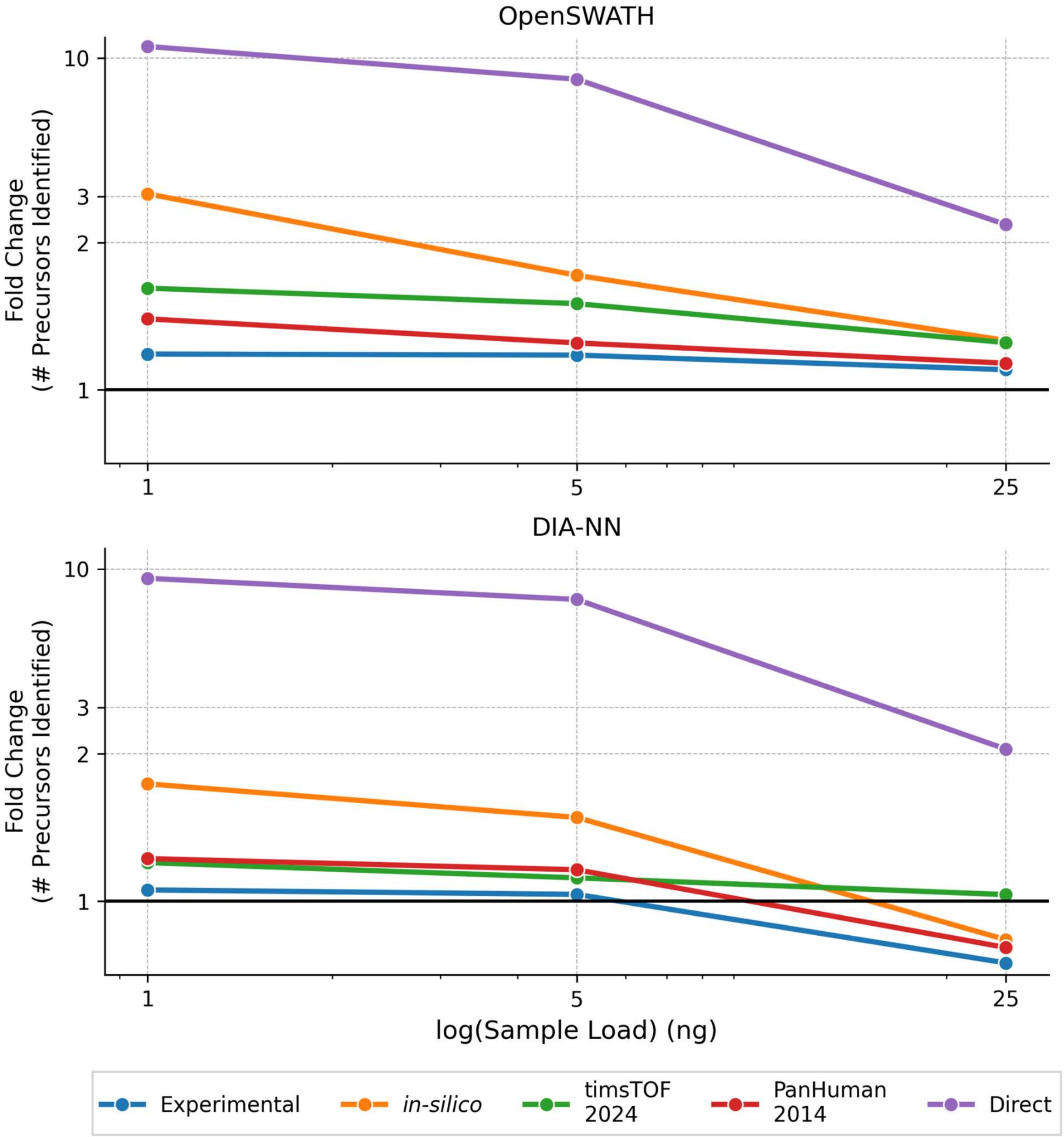


**Figure S13: Effects of reference-based library construction on precursor identifications are most pronounced in low quality libraries at the smallest sample amounts.** Fold change in peptide precursor identifications between the reference-based and the original libraries for OpenSWATH (top) and DIA-NN (bottom). x-axes are logarithmically scaled. For simplicity all reference-based libraries are constructed with the 100 ng sample as reference. Effects are most pronounced in low quality library sources at the smallest sample amounts. Across both software tools, over a 9-fold increase in peptide precursors identified is observed with the spectrum-centric construction approach at the 1 ng dilution.

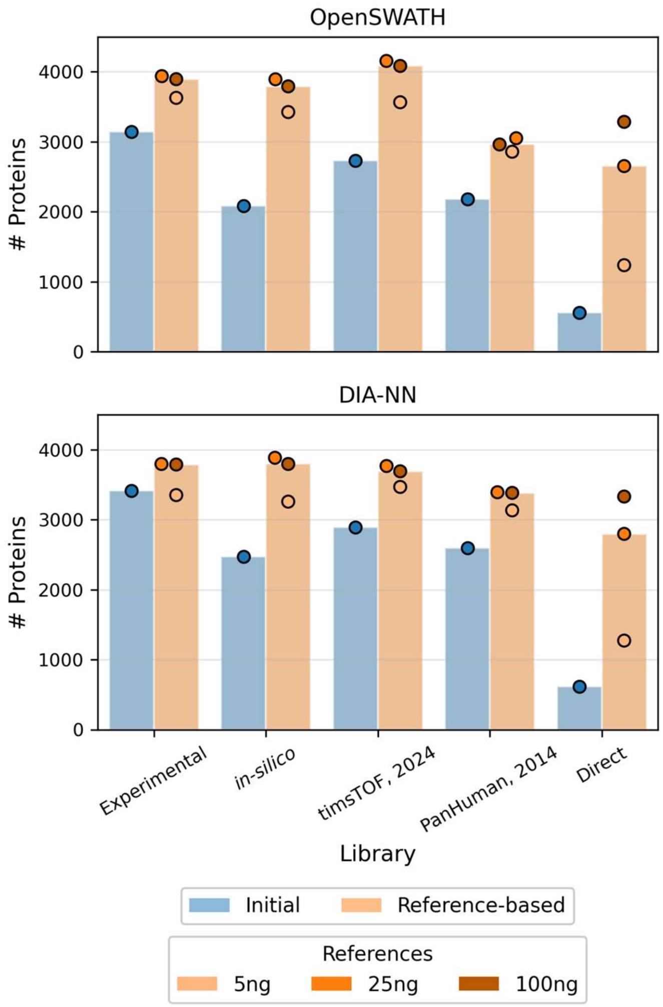


**Figure S14: At the 1ng injection, reference-based library construction consistently improves protein groups identified across libraries and software tools.** Number of protein groups identified at 1 ng sample injection across libraries for OpenSWATH (top) and DIA-NN (bottom) comparing the initial (blue) and reference-based (orange) libraries. The most prominent improvement occurs in the direct library across both software tools. Points represent the number of protein groups obtained using different reference samples for library construction, ranging from 5 ng to 100 ng, with the library bar's height representing the median identifications across the different reference dilutions. The optimal reference for maximizing protein identifications is 25 ng for peptide-centric reconstruction and 100 ng for spectrum-centric construction.

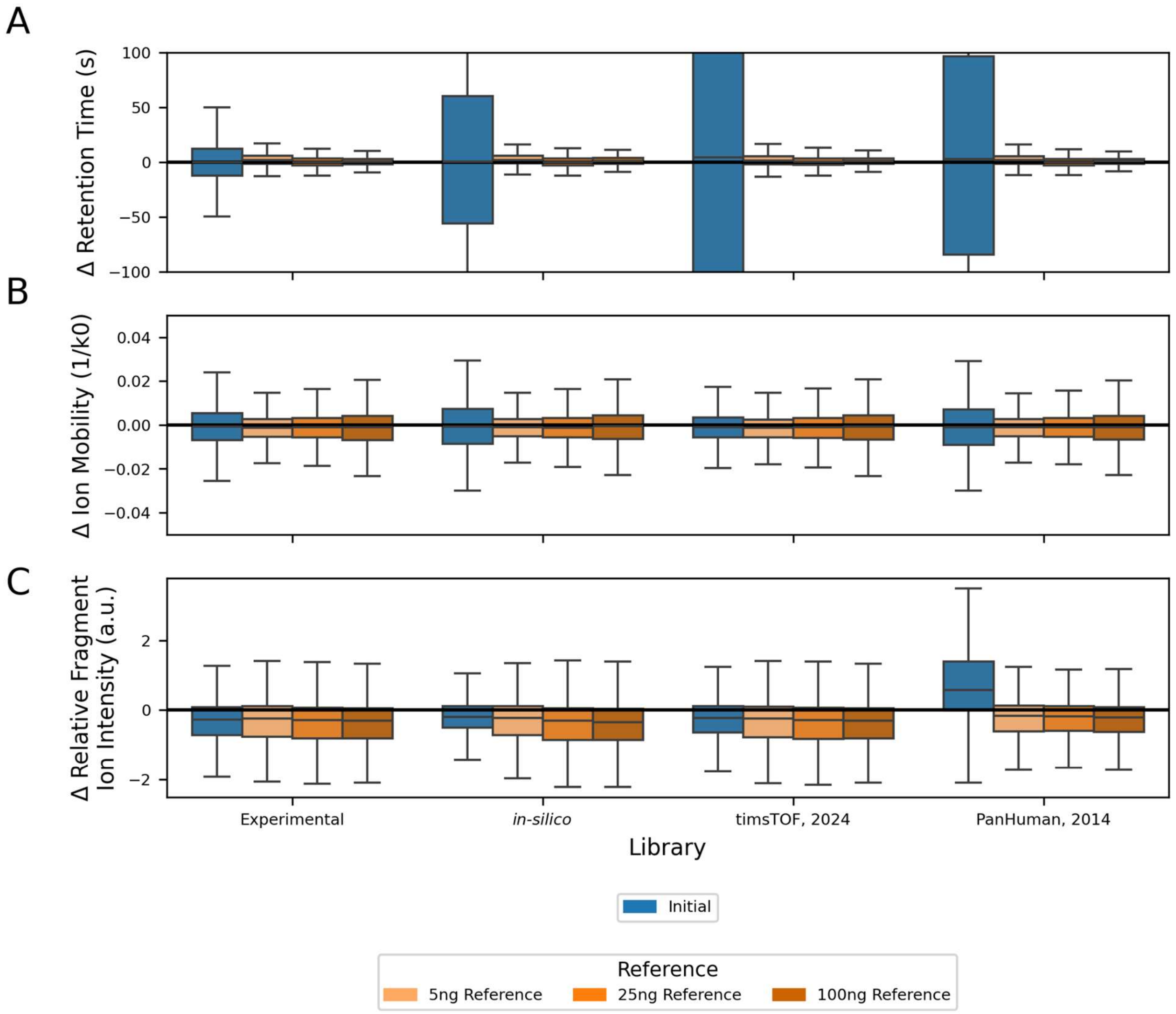


**Figure S15: Peptide-centric library reconstruction improves peptide properties precision at 1 ng sample amounts.** Comparison of peptide characteristics between the initial (blue) and reconstructed (orange) versions of the library using OpenSWATH. Multiple versions of the reconstructed library are shown (different shades of orange) using a different reference sample. Peptide characteristics refined include (A) retention time (top), (B) ion mobility (middle) and (C) relative fragment ion intensity (bottom). Tighter distributions around zero reflect more accurate library predictions relative to experimental observations. Across all libraries and reference samples, library reconstruction decreases variation in retention time and ion mobility. For retention time, the lowest variation is achieved with the largest sample amount as a reference and in ion mobility the lowest variation is achieved using the lowest sample amount as a reference. Variation in relative fragment ion intensity remains largely unchanged apart from the PanHuman, 2014 library.

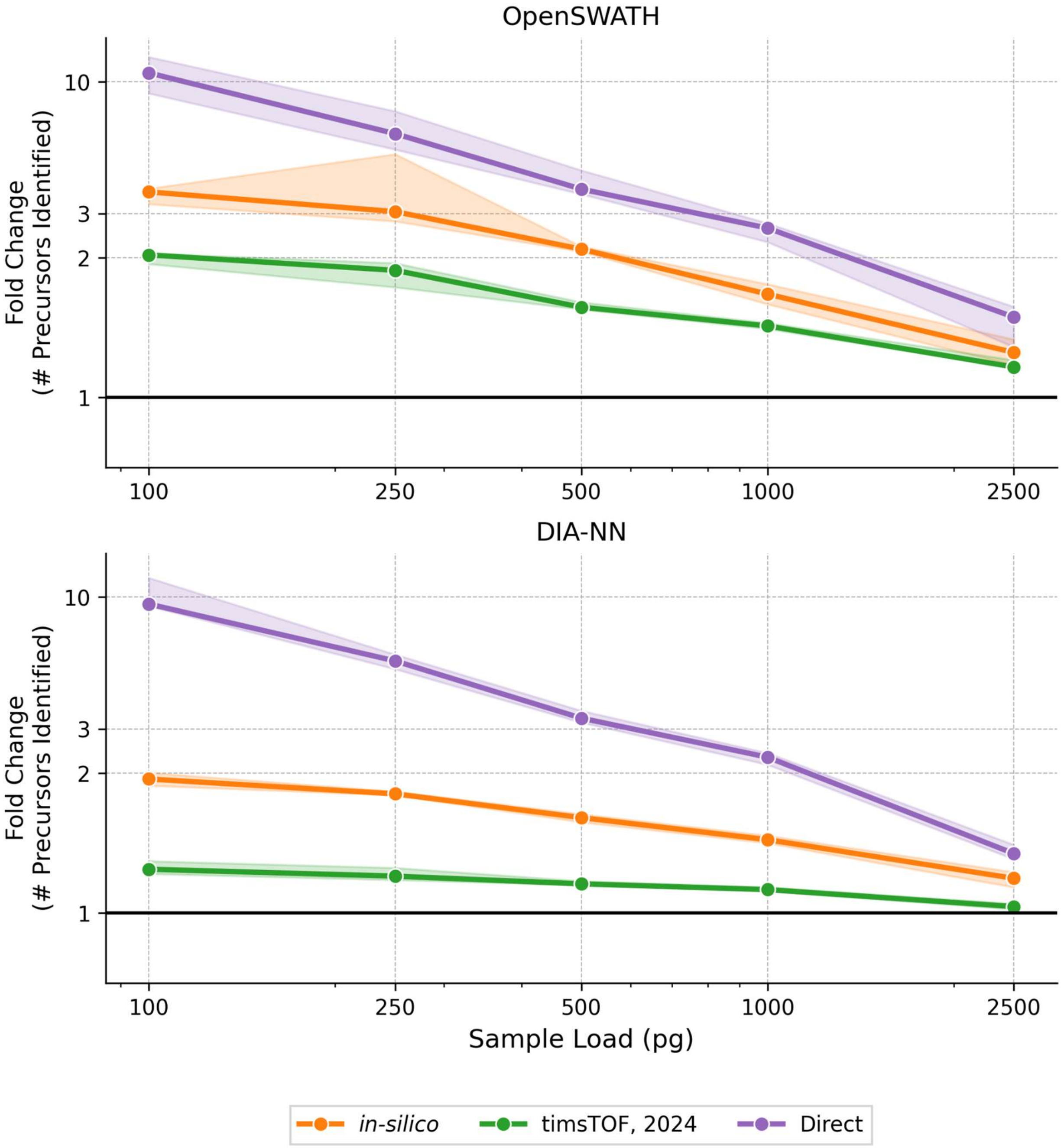


**Figure S16: Library construction effects are most pronounced at small sample amounts.** Median fold change in the number of peptide precursor identifications after reference-based library construction across the dilution series for OpenSWATH (top) and DIA-NN (bottom). Both axes are logarithmically scaled. Points are the median across the 9 replicates and the ranges indicating the 95% confidence interval across the 9 replicates. Some injections in the direct library were excluded if they failed to identify any peptide precursors. For consistency, the 5000 pg dilution load was always used as the reference. Across the libraries and software tools effects are most pronounced at the smallest sample loads. Effects are most pronounced in the spectrum-centric construction strategy across both software tools.

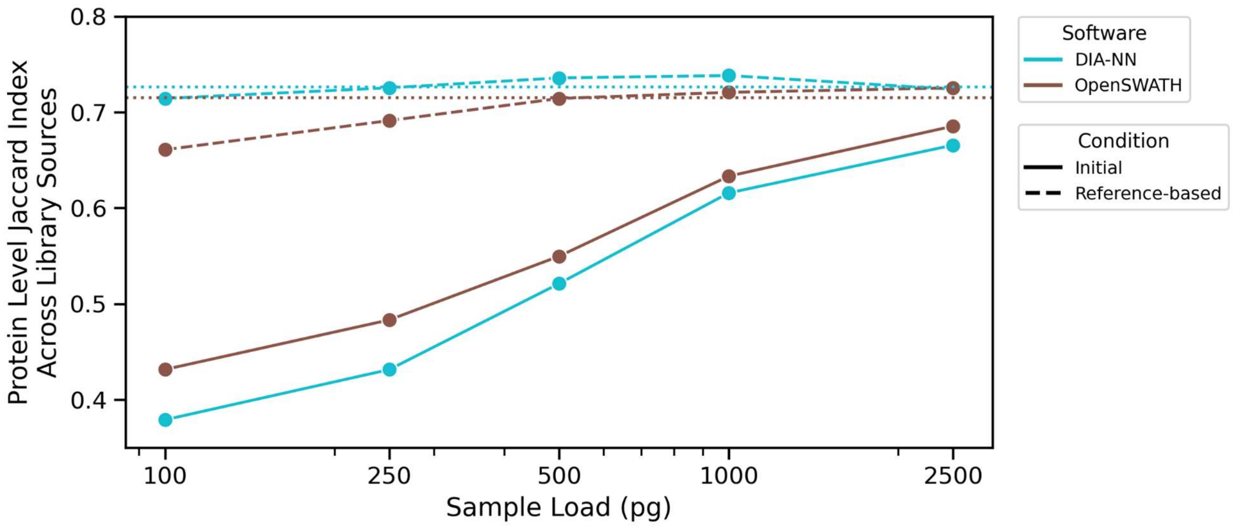


**Figure S17: Reference-based library construction decreases protein-level variation between different library sources.** Reproducibility of proteins identified across library sources for the initial (solid) and reference-based (dashed) libraries for DIA-NN (teal) and OpenSWATH (brown). x-axis is logarithmically scaled. One replicate per dilution is used. Dotted lines indicate the reproducibility achieved at the 5000 pg sample injection. Protein-level variation resulting from library source is magnified at the lowest dilution amounts. Reference-based library construction mitigates this dependency enabling consistent variation regardless of the sample amount injected.

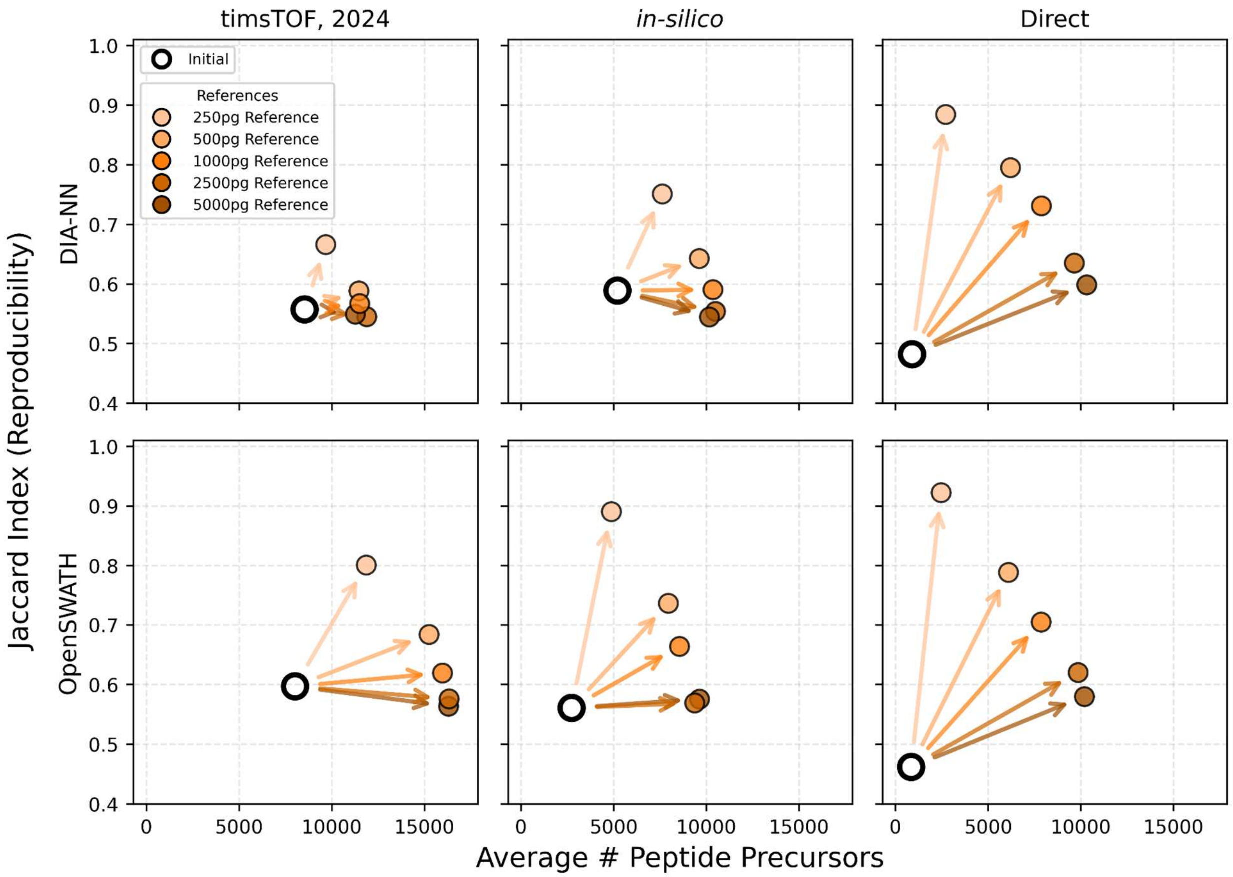


**Figure S18: Trade-off between precursor identifications and reproducibility is dependent on the reference sample.** Relationship between precursor identification and reproducibility (Jaccard index) before and after construction for the 100 pg sample for various reference samples across different library sources for DIA-NN (top) and OpenSWATH (bottom). The initial library is represented as a black circle, and the reference-based libraries are a shade of orange, with the darker shades representing a larger reference sample load. The relative influence on reproducibility and identifications is dependent on the reference sample used for library construction. Using a reference from a smaller sample amount results in prioritizing reproducibility over the number of identifications.

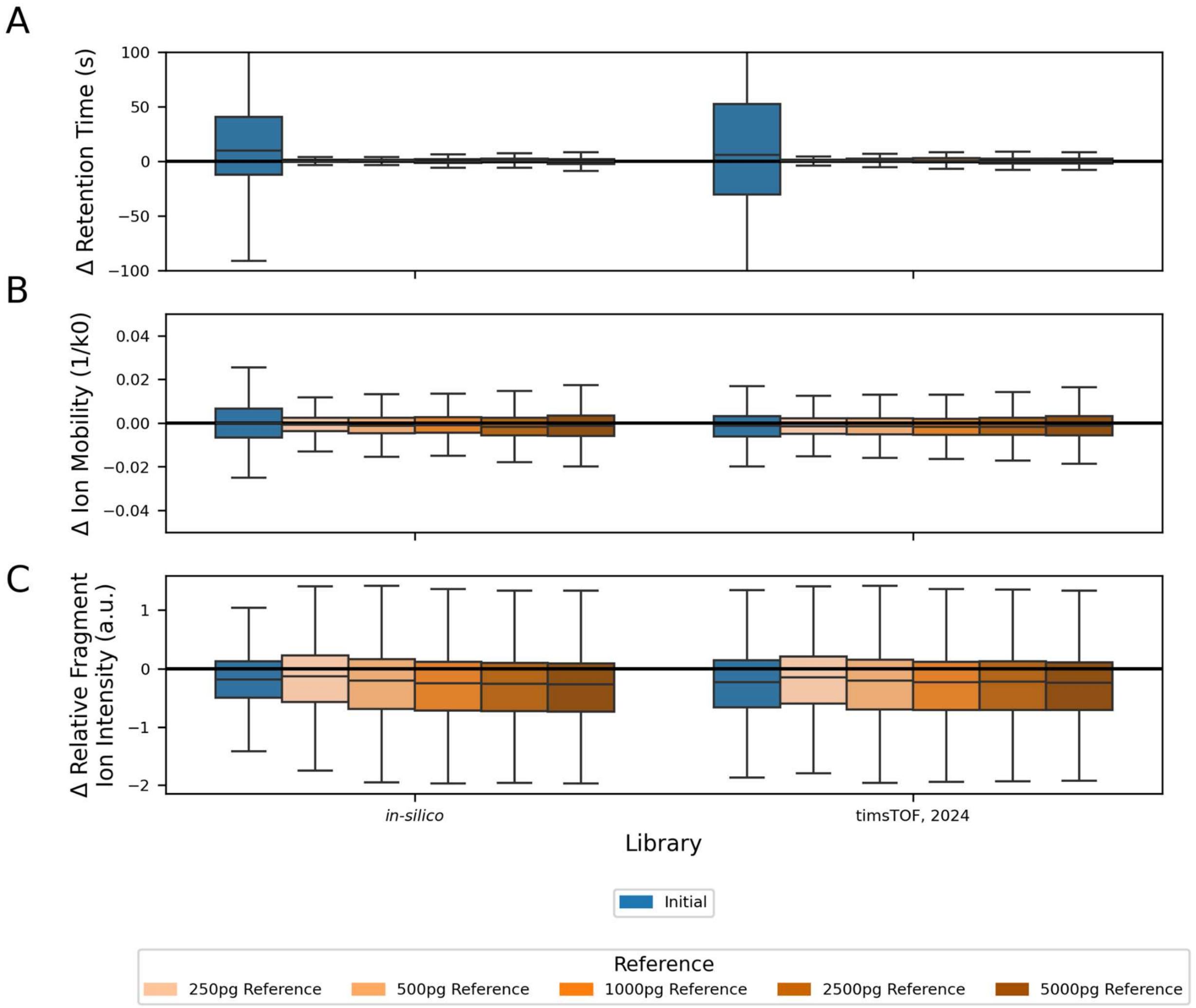


**Figure S19: Peptide-centric library reconstruction improves peptide properties precision in 100 pg sample amounts.** Comparison of peptide characteristics between the initial (blue), reconstructed (orange) versions of the library using OpenSWATH. Multiple versions of the reconstructed library are shown (different shades of orange) using a different reference sample. Peptide characteristics refined include (A) retention time (top), (B) ion mobility (middle) and (C) relative fragment ion intensity (bottom). Boxplots with low variation around zero indicate lower variation between the library and the experimentally measured value. Library reconstruction decreases variation in retention time and ion mobility across all reference samples. For retention time and ion mobility the lowest variation is achieved with the smallest sample amount as a reference. Variation in relative fragment ion intensity remains largely unchanged.

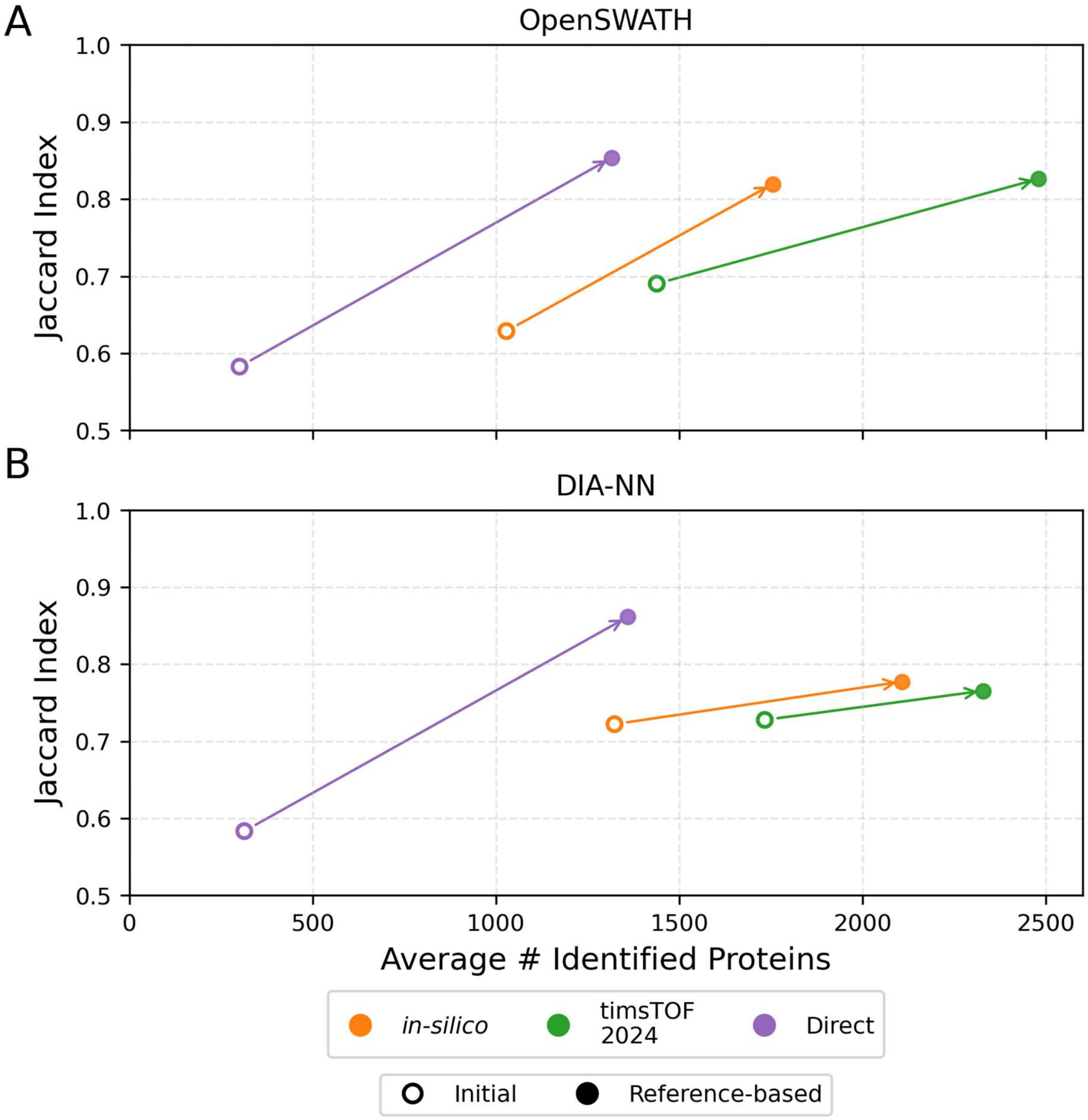


**Figure S20: Reference-based library construction increases the number of protein groups confidently detected in a 100 pg injection.** Average number of identified protein groups against reproducibility (Jaccard index) of a 100 pg sample for (A) OpenSWATH and (B) DIA-NN for the initial (blue) and reference-based (orange) libraries across different library sources. Reference-based library construction uses a 500 pg sample. Some replicates were excluded in the direct approach initial library if no confident identifications were found. The results from the initial library are represented by a hollow circle, and the results from the reference-based library are colored with a filled-in circle. Different colors represent the different libraries tested. Arrows compare the relationship between the initial and reference-based library for each library source. Arrows point to the upper right indicating improvement in protein group identifications and reproducibility. Effects are most consistently pronounced in the spectrum-centric construction workflow.

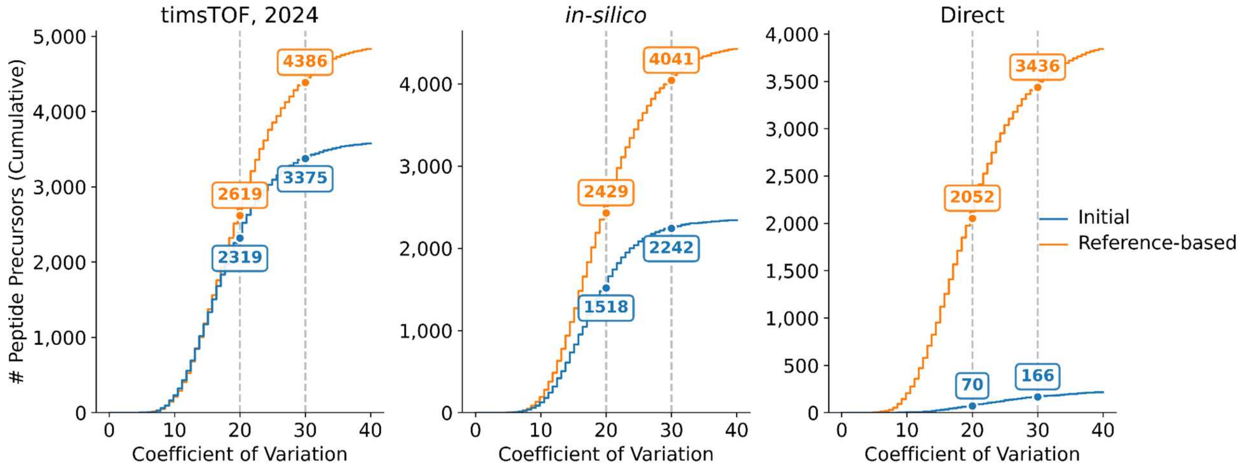


**Figure S21: Reference-based library construction increases the number of quantitatively precise peptide precursors across libraries in DIA-NN.** Cumulative distribution of commonly identified peptide precursors (1% FDR) ranked by their CV in intensity on DIA-NN results for the initial (blue) and reference-based (orange) libraries. For the direct library, replicates which did not result in any identifications were excluded. The reference-based libraries quantify on average 1050 and 2000 more peptides with an intensity CV less than 20% and 30% respectively across all library sources.

**Table S1: GPF Acquisition Scheme**

| Injection Number | m/z Window | Mobility Range |
|---|---|---|
| 1 | 400 - 405 | 0.60 - 0.98 |
| 1 | 404 - 409 | 0.60 - 1.03 |
| 1 | 409 - 414 | 0.60 - 1.08 |
| 1 | 414 - 419 | 0.60 - 1.13 |
| 1 | 419 - 424 | 0.60 - 1.18 |
| 1 | 424 - 429 | 0.60 - 1.23 |
| 1 | 429 - 434 | 0.60 - 1.28 |
| 1 | 434 - 439 | 0.60 - 1.34 |
| 1 | 439 - 444 | 0.60 - 1.39 |
| 1 | 444 - 449 | 0.60 - 1.44 |
| 1 | 449 - 454 | 0.60 - 1.49 |
| 1 | 454 - 459 | 0.60 - 1.54 |
| 1 | 459 - 464 | 0.60 - 1.59 |
| 2 | 460 - 465 | 0.60 - 0.94 |
| 2 | 464 - 469 | 0.60 - 0.99 |
| 2 | 469 - 474 | 0.60 - 1.04 |
| 2 | 474 - 479 | 0.60 - 1.09 |
| 2 | 479 - 484 | 0.60 - 1.14 |
| 2 | 484 - 489 | 0.60 - 1.19 |
| 2 | 489 - 494 | 0.60 - 1.25 |
| 2 | 494 - 499 | 0.60 - 1.30 |
| 2 | 499 - 504 | 0.60 - 1.35 |
| 2 | 504 - 509 | 0.60 - 1.40 |
| 2 | 509 - 514 | 0.60 - 1.45 |
| 2 | 514 - 519 | 0.60 - 1.50 |
| 2 | 519 - 524 | 0.60 - 1.55 |
| 3 | 520 - 525 | 0.60 - 0.98 |
| 3 | 524 - 529 | 0.60 - 1.04 |
| 3 | 529 - 534 | 0.60 - 1.09 |
| 3 | 534 - 539 | 0.60 - 1.14 |
| 3 | 539 - 544 | 0.60 - 1.19 |
| 3 | 544 - 549 | 0.60 - 1.24 |
| 3 | 549 - 554 | 0.60 - 1.29 |
| 3 | 554 - 559 | 0.60 - 1.34 |
| 3 | 559 - 564 | 0.60 - 1.40 |
| 3 | 564 - 569 | 0.60 - 1.45 |
| 3 | 569 - 574 | 0.61 - 1.50 |
| 3 | 574 - 579 | 0.66 - 1.55 |
| 3 | 579 - 584 | 0.71 - 1.60 |
| 4 | 580 - 585 | 0.60 - 1.06 |
| 4 | 584 - 589 | 0.60 - 1.11 |
| 4 | 589 - 594 | 0.60 - 1.16 |
| 4 | 594 - 599 | 0.60 - 1.21 |
| 4 | 599 - 604 | 0.60 - 1.26 |

| | | |
|---|---|---|
| 4 | 604 - 609 | 0.60 - 1.31 |
| 4 | 609 - 614 | 0.60 - 1.36 |
| 4 | 614 - 619 | 0.60 - 1.41 |
| 4 | 619 - 624 | 0.60 - 1.47 |
| 4 | 624 - 629 | 0.63 - 1.52 |
| 4 | 629 - 634 | 0.68 - 1.57 |
| 4 | 634 - 639 | 0.73 - 1.60 |
| 4 | 639 - 644 | 0.78 - 1.60 |
| 5 | 640 - 645 | 0.60 - 1.09 |
| 5 | 644 - 649 | 0.60 - 1.14 |
| 5 | 649 - 654 | 0.60 - 1.19 |
| 5 | 654 - 659 | 0.60 - 1.24 |
| 5 | 659 - 664 | 0.60 - 1.30 |
| 5 | 664 - 669 | 0.60 - 1.35 |
| 5 | 669 - 674 | 0.60 - 1.40 |
| 5 | 674 - 679 | 0.60 - 1.45 |
| 5 | 679 - 684 | 0.62 - 1.50 |
| 5 | 684 - 689 | 0.67 - 1.55 |
| 5 | 689 - 694 | 0.72 - 1.60 |
| 5 | 694 - 699 | 0.77 - 1.60 |
| 5 | 699 - 704 | 0.82 - 1.60 |
| 6 | 700 - 705 | 0.60 - 1.13 |
| 6 | 704 - 709 | 0.60 - 1.18 |
| 6 | 709 - 714 | 0.60 - 1.23 |
| 6 | 714 - 719 | 0.60 - 1.28 |
| 6 | 719 - 724 | 0.60 - 1.34 |
| 6 | 724 - 729 | 0.60 - 1.39 |
| 6 | 729 - 734 | 0.60 - 1.44 |
| 6 | 734 - 739 | 0.60 - 1.49 |
| 6 | 739 - 744 | 0.65 - 1.54 |
| 6 | 744 - 749 | 0.71 - 1.59 |
| 6 | 749 - 754 | 0.76 - 1.60 |
| 6 | 754 - 759 | 0.81 - 1.60 |
| 6 | 759 - 764 | 0.86 - 1.60 |
| 7 | 760 - 765 | 0.60 - 0.98 |
| 7 | 764 - 769 | 0.60 - 1.04 |
| 7 | 769 - 774 | 0.60 - 1.09 |
| 7 | 774 - 779 | 0.60 - 1.14 |
| 7 | 779 - 784 | 0.60 - 1.19 |
| 7 | 784 - 789 | 0.60 - 1.24 |
| 7 | 789 - 794 | 0.60 - 1.29 |
| 7 | 794 - 799 | 0.60 - 1.34 |
| 7 | 799 - 804 | 0.65 - 1.40 |
| 7 | 804 - 809 | 0.71 - 1.45 |
| 7 | 809 - 814 | 0.76 - 1.50 |
| 7 | 814 - 819 | 0.81 - 1.55 |

| | | |
|---|---|---|
| 7 | 819 - 824 | 0.86 - 1.60 |
| 8 | 820 - 825 | 0.60 - 1.26 |
| 8 | 824 - 829 | 0.60 - 1.31 |
| 8 | 829 - 834 | 0.60 - 1.36 |
| 8 | 834 - 839 | 0.60 - 1.42 |
| 8 | 839 - 844 | 0.60 - 1.47 |
| 8 | 844 - 849 | 0.60 - 1.52 |
| 8 | 849 - 854 | 0.60 - 1.57 |
| 8 | 854 - 859 | 0.65 - 1.60 |
| 8 | 859 - 864 | 0.70 - 1.60 |
| 8 | 864 - 869 | 0.76 - 1.60 |
| 8 | 869 - 874 | 0.81 - 1.60 |
| 8 | 874 - 879 | 0.86 - 1.60 |
| 8 | 879 - 884 | 0.91 - 1.60 |
| 9 | 880 - 885 | 0.60 - 1.29 |
| 9 | 884 - 889 | 0.60 - 1.34 |
| 9 | 889 - 894 | 0.60 - 1.39 |
| 9 | 894 - 899 | 0.60 - 1.44 |
| 9 | 899 - 904 | 0.60 - 1.49 |
| 9 | 904 - 909 | 0.62 - 1.54 |
| 9 | 909 - 914 | 0.67 - 1.59 |
| 9 | 914 - 919 | 0.72 - 1.60 |
| 9 | 919 - 924 | 0.78 - 1.60 |
| 9 | 924 - 929 | 0.83 - 1.60 |
| 9 | 929 - 934 | 0.88 - 1.60 |
| 9 | 934 - 939 | 0.93 - 1.60 |
| 9 | 939 - 944 | 0.98 - 1.60 |
| 10 | 940 - 945 | 0.60 - 1.34 |
| 10 | 944 - 949 | 0.60 - 1.39 |
| 10 | 949 - 954 | 0.60 - 1.44 |
| 10 | 954 - 959 | 0.60 - 1.50 |
| 10 | 959 - 964 | 0.60 - 1.55 |
| 10 | 964 - 969 | 0.60 - 1.60 |
| 10 | 969 - 974 | 0.60 - 1.60 |
| 10 | 974 - 979 | 0.61 - 1.60 |
| 10 | 979 - 984 | 0.66 - 1.60 |
| 10 | 984 - 989 | 0.71 - 1.60 |
| 10 | 989 - 994 | 0.76 - 1.60 |
| 10 | 994 - 999 | 0.81 - 1.60 |
| 10 | 999 - 1004 | 0.86 - 1.60 |
| 11 | 1000 - 1005 | 0.60 - 1.40 |
| 11 | 1004 - 1009 | 0.60 - 1.45 |
| 11 | 1009 - 1014 | 0.60 - 1.50 |
| 11 | 1014 - 1019 | 0.60 - 1.55 |
| 11 | 1019 - 1024 | 0.60 - 1.60 |
| 11 | 1024 - 1029 | 0.64 - 1.60 |

| | | |
|---|---|---|
| 11 | 1029 - 1034 | 0.69 - 1.60 |
| 11 | 1034 - 1039 | 0.74 - 1.60 |
| 11 | 1039 - 1044 | 0.79 - 1.60 |
| 11 | 1044 - 1049 | 0.85 - 1.60 |
| 11 | 1049 - 1054 | 0.90 - 1.60 |
| 11 | 1054 - 1059 | 0.95 - 1.60 |
| 11 | 1059 - 1064 | 1.00 - 1.60 |
| 12 | 1060 - 1065 | 0.60 - 1.42 |
| 12 | 1064 - 1069 | 0.60 - 1.47 |
| 12 | 1069 - 1074 | 0.60 - 1.52 |
| 12 | 1074 - 1079 | 0.60 - 1.58 |
| 12 | 1079 - 1084 | 0.60 - 1.60 |
| 12 | 1084 - 1089 | 0.64 - 1.60 |
| 12 | 1089 - 1094 | 0.69 - 1.60 |
| 12 | 1094 - 1099 | 0.74 - 1.60 |
| 12 | 1099 - 1104 | 0.79 - 1.60 |
| 12 | 1104 - 1109 | 0.85 - 1.60 |
| 12 | 1109 - 1114 | 0.90 - 1.60 |
| 12 | 1114 - 1119 | 0.95 - 1.60 |
| 12 | 1119 - 1124 | 1.00 - 1.60 |
| 13 | 1120 - 1125 | 0.60 - 1.41 |
| 13 | 1124 - 1129 | 0.60 - 1.45 |
| 13 | 1129 - 1134 | 0.60 - 1.49 |
| 13 | 1134 - 1139 | 0.60 - 1.53 |
| 13 | 1139 - 1144 | 0.60 - 1.57 |
| 13 | 1144 - 1149 | 0.60 - 1.60 |
| 13 | 1149 - 1154 | 0.60 - 1.60 |
| 13 | 1154 - 1159 | 0.64 - 1.60 |
| 13 | 1159 - 1164 | 0.68 - 1.60 |
| 13 | 1164 - 1169 | 0.73 - 1.60 |
| 13 | 1169 - 1174 | 0.77 - 1.60 |
| 13 | 1174 - 1179 | 0.81 - 1.60 |
| 13 | 1179 - 1184 | 0.85 - 1.60 |
| 13 | 1184 - 1189 | 0.89 - 1.60 |
| 13 | 1189 - 1194 | 0.93 - 1.60 |
| 13 | 1194 - 1199 | 0.97 - 1.60 |

**Table S2: Acquisition Scheme for single-cell equivalent injections**

| m/z Window | Mobility Range |
|---|---|
| 400-435 | 0.64-0.84 |
| 434-471 | 0.64-0.89 |
| 470-501 | 0.64-0.91 |
| 500-530 | 0.64-0.93 |
| 529-558 | 0.64-0.96 |
| 557-587 | 0.84-0.97 |
| 586-618 | 0.89-0.99 |
| 617-650 | 0.91-1.01 |
| 649-684 | 0.93-1.05 |
| 683-723 | 0.96-1.11 |
| 722-767 | 0.97-1.45 |
| 766-819 | 0.99-1.45 |
| 818-883 | 1.01-1.45 |
| 882-956 | 1.05-1.45 |
| 955-1000 | 1.11-1.45 |
